# Combinatorial control of type IVa pili formation by the four polarized regulators MglA, SgmX, FrzS and SopA

**DOI:** 10.1101/2024.03.11.584430

**Authors:** Michel Oklitschek, Luís António Menezes Carreira, Memduha Muratoğlu, Lotte Søgaard-Andersen, Anke Treuner-Lange

## Abstract

Type IVa pili (T4aP) are widespread and enable bacteria to translocate across surfaces. T4aP engage in cycles of extension, surface adhesion and retraction, thereby pulling cells forward. Accordingly, the number and localization of T4aP are critical to efficient translocation. Here, we address how T4aP formation is regulated in *Myxococcus xanthus*, which translocates with a well-defined leading and lagging cell pole using T4aP at the leading pole. This localization is orchestrated by the small GTPase MglA and its downstream effector SgmX that both localize at the leading pole and recruit the PilB extension ATPase to the T4aP machinery at this pole. Here, we identify the previously uncharacterized protein SopA and show that it interacts directly with SgmX, localizes at the leading pole, stimulates polar localization of PilB, and is important for T4aP formation. We corroborate that MglA also recruits FrzS to the leading pole, and that FrzS stimulates SgmX recruitment. In addition, FrzS and SgmX separately recruit SopA. Precise quantification of T4aP formation and T4aP-dependent motility in various mutants support a model whereby the main pathway for stimulating T4aP formation is the MglA/SgmX pathway. FrzS stimulates this pathway by recruiting SgmX and SopA. SopA stimulates the MglA/SgmX pathway by stimulating the function of SgmX, likely by promoting the SgmX-dependent recruitment of PilB. The architecture of the MglA/SgmX/FrzS/SopA protein interaction network for orchestrating T4aP formation allows for combinatorial regulation of T4aP levels at the leading cell pole resulting in discrete levels of T4aP-dependent motility.

## Introduction

Bacterial motility is important for colonization of environmental niches, interactions with host cells, virulence, biofilm formation and fitness by directing cells towards nutrients and away from toxins and predators (1). For translocation on solid surfaces, bacteria most commonly use type IVa pili (T4aP), long thin filaments that are also important for adhesion to host cells and abiotic surfaces, biofilm formation, virulence, predation, protein secretion, DNA uptake and surface sensing (2). T4aP undergo cycles of extension, surface adhesion and retraction (3–5). During these cycles, retractions generate a force up to 150pN that is sufficient to pull a cell forward (3, 5, 6). Efficient T4aP-dependent translocation depends on the number and cellular localization of T4aP (7, 8).

The T4aP extension/adhesion/retraction cycles are powered by the highly conserved T4aP machine (T4aPM) (2). In Gram-negative bacteria, this nanomachine is composed of 15 highly conserved proteins and spans from the outer membrane (OM) across the periplasm and inner membrane (IM) to the cytoplasm (9–11) (Fig. S1A). The hexameric PilB and PilT ATPases (12–15) associate with the cytoplasmic base of the core T4aPM in a mutually exclusive fashion to power T4aP extension and retraction, respectively (10). With the exception of PilT, all T4aPM proteins are important for T4aP extension, while PilT is only important for retraction (2). The T4aP is composed of thousands of copies of the major pilin subunit and contains a tip complex composed of minor pilins and the PilY1 adhesin (11, 16–18). During extensions, major pilins are extracted from the IM and inserted at the T4aP base (4, 19, 20); during retractions, this process is inverted and major pilin subunits removed from the T4aP base and reinserted into the IM (4, 21). While the highly conserved T4aPM constitutes the basis for the extension/adhesion/retraction cycles, much-less conserved regulatory proteins determine where and how many T4aP are formed (7, 18, 22–29). However, their mechanism of action is poorly understood. Here, we address the regulation of T4aP formation in *Myxococcus xanthus*, a predatory soil bacterium with a social lifestyle and a model organism for understanding T4aPM function and regulation.

The rod-shaped *M. xanthus* cells move across surfaces in the direction of their long axis using two motility systems, one for gliding and one for T4aP-dependent motility (30, 31). Motility is important for the social behaviors of *M. xanthus* including predation and formation of swarming colonies in the presence and spore-filled fruiting bodies in the absence of nutrients (30–32). The T4aPM core is present at both cell poles (11, 33–37). However, T4aP only assemble at one pole at a time (38, 39). This localization enables *M. xanthus* cells to move unidirectionally with a piliated leading and a non-piliated lagging cell pole (7, 39) and is essential for efficient translocation across surfaces (7). Consistent with the unipolar T4aP formation, the PilB extension ATPase localizes to the leading cell pole, while PilT predominantly localizes to the lagging pole and only occasionally localizes to the leading pole stimulating retractions (34). In response to signaling by the Frz chemosensory system, *M. xanthus* cells reverse their direction of translocation (40) and after a reversal, T4aP assemble at the new leading pole (39); in parallel, PilB and PilT switch polarity (34).

The activity of the T4aPM in *M. xanthus* is regulated by the polarity module (41–43). The output of this module is generated by the small Ras-like GTPase MglA, which is a nucleotide-dependent molecular switch that is inactive in the GDP-bound and active in the GTP-bound state (44, 45). In its GTP-bound state MglA localizes to and defines the leading cell pole (44, 45) (Fig. S1B). At this pole, MglA interacts with effectors to stimulate the T4aPM resulting in T4aP formation (7, 46) and is essential for T4aP-dependent motility (47, 48). The remaining five proteins regulate the nucleotide-bound state and localization of MglA by acting as a guanine nucleotide exchange factor (GEF) in case of the RomR/RomX complex (49) or as a GTPase activating protein (GAP) in case of the MglB/RomY complex (44, 45, 50). MglA and the RomR/RomX and MglB/RomY complexes together with the MglC protein interact to bring about their asymmetric polar localization (43, 51) (Fig. S1B). During the Frz-induced reversals, these six proteins switch polarity, thereby enabling the activation of the T4aPM at the new leading cell pole (43–45, 49, 50, 52–54).

At the leading pole, MglA directly interacts with and recruits SgmX, a protein containing 14 tetratricopeptide repeats (TPR) (7, 46), and has also been suggested to interact directly with FrzS (55), which is also important for T4aP-dependent motility (56, 57). FrzS also interacts directly with SgmX and stimulates the recruitment of SgmX to the leading pole (58). SgmX, in turn, brings about PilB localization at the leading pole by an unknown mechanism and is essential for T4aP formation and, consequently, also for T4aP-dependent motility (7). Based on these observations, it has been suggested that SgmX stimulates T4aP formation by enabling PilB interaction with the base of the T4aPM (7).

Here, to increase our understanding of how T4aP formation is regulated in *M. xanthus*, we searched for putative SgmX interaction partners. We identify the previously uncharacterized protein MXAN_0371 (reannotated to MXAN_RS01825 in the NCBI Reference Sequence NC_008095.1; henceforth <u>S</u>timulation of <u>p</u>ili formation <u>p</u>rotein <u>A</u>, SopA) and demonstrate that SopA interacts directly with SgmX, localizes at the leading pole, stimulates polar PilB localization, and is important but not essential for T4aP formation and T4aP-dependent motility. We confirm that MglA is important but not essential for FrzS polar localization and that FrzS interacts directly with SgmX, thereby stimulating the polar recruitment of SgmX. In doing so, FrzS indirectly stimulates PilB polar localization, T4aP formation and T4aP-dependent motility. Additionally, SgmX and FrzS can separately recruit SopA to the leading pole. Altogether, our data support a model whereby MglA, SgmX, FrzS and SopA interact to establish a protein interaction network that allows for combinatorial regulation of T4aP formation at the leading cell pole resulting in discrete levels of T4aP-dependent motility.

## Results

### SopA is important for T4aP-dependent motility

We previously identified RomX and RomY using a phylogenomic approach in which we searched for proteins that co-occur with MglA, MglB and RomR of the polarity module (49, 50). Therefore, to identify proteins that could interact with SgmX, we searched the STRING database (59) for proteins that co-occur with SgmX, resulting in the identification of 10 proteins (Table S1). With the exception of MXAN_5763-_5765 (reannotated to MXAN_RS27935, MXAN_RS27940 and MXAN_RS27945 in the NCBI Reference Sequence NC_008095.1), which are encoded downstream of *sgmX* (7, 28), and deletion of which has no impact on T4aP-dependent motility (60), none of these proteins have previously been analyzed. Three of the remaining seven proteins are predicted to have enzymatic activity and were not considered further. The hypothetical protein SopA (MXAN_0371/MXAN_RS01825) and the TPR domain protein MXAN_6595 (reannotated to MXAN_RS24110) are both highly conserved in Myxococcales with fully sequenced genomes (Fig. S2A), while the PATAN domain proteins MXAN_3211 (reannotated to MXAN_RS15550) and MXAN_4965 (reannotated to MXAN_RS24110) are less conserved. From here on, we focused on SopA.

Based on sequence analysis, SopA is a 405 amino acid residue cytoplasmic protein and homologs were only identified in Myxococcales. SopA homologs share conserved N- and C-terminal regions, which do not match characterized domain models (Fig. 1A, Fig. S3). While the *sopA* locus is conserved in related Myxococcales, none of the genes flanking *sopA* have been implicated in motility (Fig. S2B). Based on RNAseq and cappableseq analyses (61), *sopA* is not encoded in an operon (Fig. 1A).

**Figure 1.**
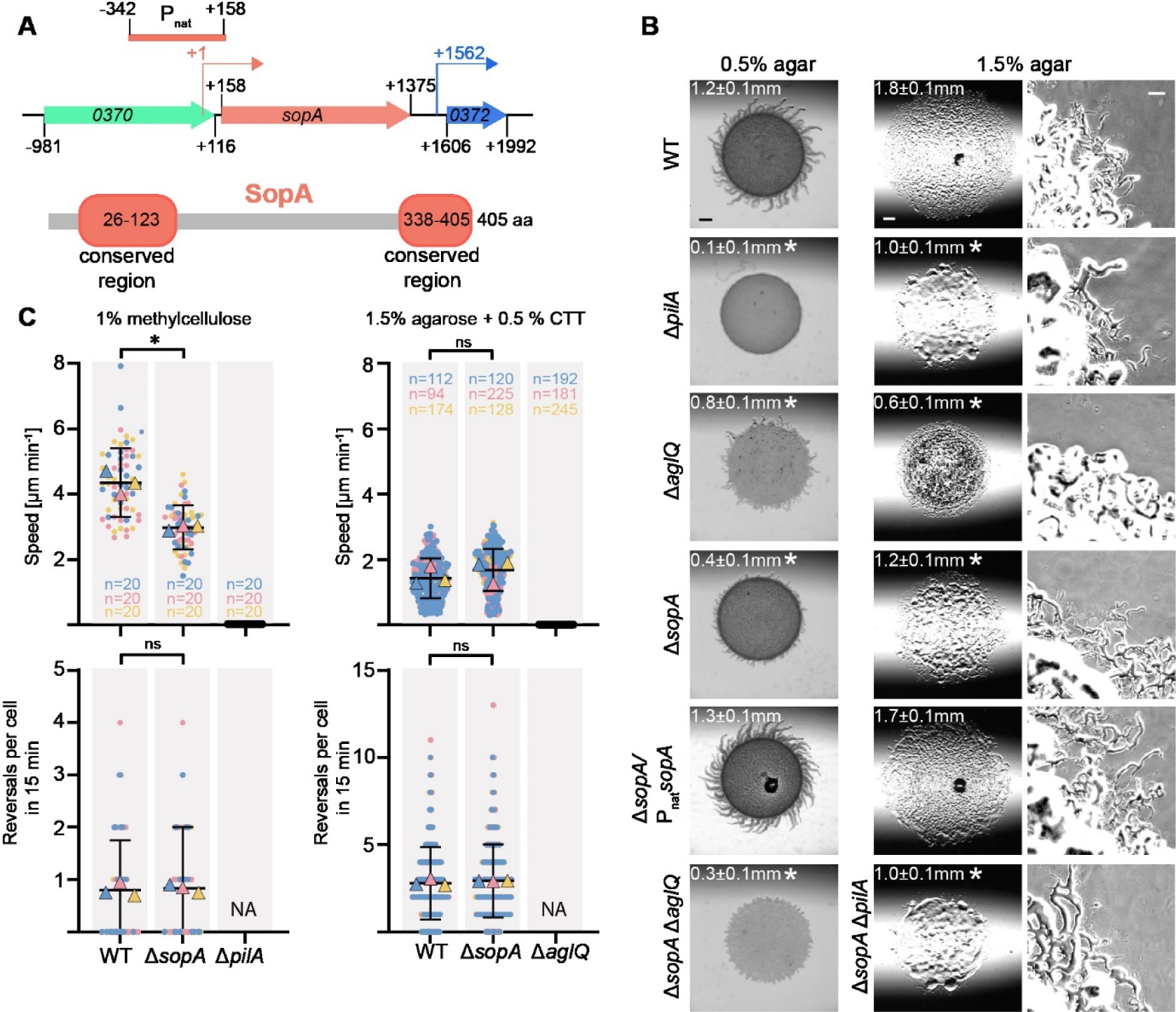
SopA is important for T4aP-dependent motility. **A.** *sopA* locus and SopA domain architecture. Upper panel, *sopA* locus; numbers in arrows, MXAN locus tags (in the NCBI Reference Sequence NC_008095.1, MXAN_0370 and MXAN_0372 are reannotated as MXAN_RS01820 and MXAN_RS01830, respectively); numbers in black indicate the first and last nucleotide in start and stop codons, respectively relative to +1, the transcriptional start site of *sopA* (61). Kinked arrows, transcriptional start sites. Light red bar labelled P_nat_ indicate the 500 bp fragment upstream of the *sopA* start codon used for ectopic expression of *sopA*. Lower panel, conserved regions of SopA homologs are indicated in light red using the coordinates of SopA of *M. xanthus*. **B.** SopA is important for T4aP-dependent motility in population-based assay. T4aP-dependent motility and gliding were analyzed on 0.5% and 1.5% agar supplemented with 0.5% CTT, respectively. Numbers indicate the colony expansion in 24 h as mean ± standard deviation (SD) (*n*=3 biological replicates). * *P*<0.05, two-tailed Student’s *t-*test for samples with equal variances. In the complementation strain, *sopA* was expressed from its native promoter from a plasmid integrated in a single copy at the Mx8 *attB* site. Scale bars, 1 mm (left, middle), 100 µm (right). **C.** SopA is important for T4aP-dependent motility in single cell-based motility assay. T4aP-dependent motility was measured for cells on a polystyrene surface covered with 1% methylcellulose and gliding on 1.5% agar supplemented with 0.5% CTT. Individual data points from three biological replicates indicated in three different colors and with the number of cells per replicate indicated in the corresponding colors. The mean is shown for each experiment and the mean for all experiments ± SD is shown in black. * *P*<0.05, two-tailed Student’s *t-*test for samples with equal variances, ns, not significant, NA, not applicable because cells are non-motile.

To characterize a potential function of SopA in motility, we generated a *sopA* in-frame deletion mutant (Δ*sopA*) in the DK1622 wild-type (WT) strain and analyzed the motility characteristics of Δ*sopA* cells in population-based assays. In motility assays on 0.5% agar supplemented with 0.5% casitone broth (CTT), which is most favorable for T4aP-dependent motility (62), WT displayed the long flares at the edge of colonies characteristic of T4aP-dependent motility, while the Δ*pilA* mutant, which lacks the major pilin of T4aP (63) and served as a negative control, generated smooth colony edges without flares (Fig. 1B). The Δ*sopA* mutant formed significantly shorter flares than WT and was significantly reduced in colony expansion (Fig. 1B). This motility defect was complemented by the ectopic expression of *sopA* from its native promoter from a plasmid integrated in a single copy at the Mx8 *attB* site (Fig. 1A-B). Because the Δ*aglQ* mutant, which has a defect in gliding motility due to the lack of a component of the Agl/Glt machinery for gliding (64, 65), also exhibited reduced flare formation on 0.5% agar, we compared flare formation and colony expansion of the Δ*aglQ* mutant and the Δ*sopA*Δ*aglQ* double mutant. The Δ*sopA*Δ*aglQ* double mutant exhibited significantly shorter flares and reduced in colony expansion compared to the Δ*aglQ* mutant (Fig. 1B), documenting that the Δ*sopA* mutation causes a defect in T4aP-dependent motility. On 1.5% agar supplemented with 0.5% CTT, which is most favorable for gliding (62), WT displayed single cells at the edge of the colony, which was not the case for the Δ*aglQ* mutant, which served as a negative control (Fig. 1B). The Δ*sopA* mutant also exhibited single cells at the colony edge but was significantly reduced in colony expansion, and this motility defect was complemented by the ectopic expression of *sopA* (Fig. 1B). Because the Δ*pilA* mutant, while still displaying single cells at the colony edge, also had reduced colony expansion on 1.5% agar, we compared its motility characteristics with those of the Δ*sopA*Δ*pilA* double mutant. These two strains had the same colony expansion and both had single cells at the colony edge (Fig. 1B). Thus, SopA is not important for gliding motility.

A motility defect in the population-based assay can be caused by a *bona fide* motility defect or by an altered reversal frequency. To distinguish between these two possibilities, we analyzed the single cell behavior of Δ*sopA* cells. In the single cell assay for T4aP-dependent motility, cells of the Δ*sopA* mutant displayed a significantly reduced speed compared to WT, while the reversal frequency was unaffected (Fig. 1C). In the single cell assay for gliding, cells of the Δ*sopA* mutant displayed the same speed and reversal frequency as WT (Fig. 1C).

Based on these motility assays, we conclude that SopA is important but not essential for T4aP-dependent motility and is not important for gliding motility. Moreover, lack of SopA does not interfere with proper reversals. By comparison, SgmX is essential for T4aP-dependent motility (7, 46).

### SopA is important for T4aP extension

To address the mechanism underlying the defect in T4aP-dependent motility in the Δ*sopA* mutant, we examined whether this mutant assembles T4aP using an assay in which T4aP are sheared-off the cell surface followed by quantification of PilA levels by immunoblotting. PilA was still present in the sheared T4aP fraction from the Δ*sopA* mutant but at a significantly reduced level compared to WT while the total cellular level of PilA was as in WT (Fig. 2A). This defect in T4aP formation was corrected in the complementation strain in which *sopA* was ectopically expressed (Fig. 2A).

**Figure 2.**
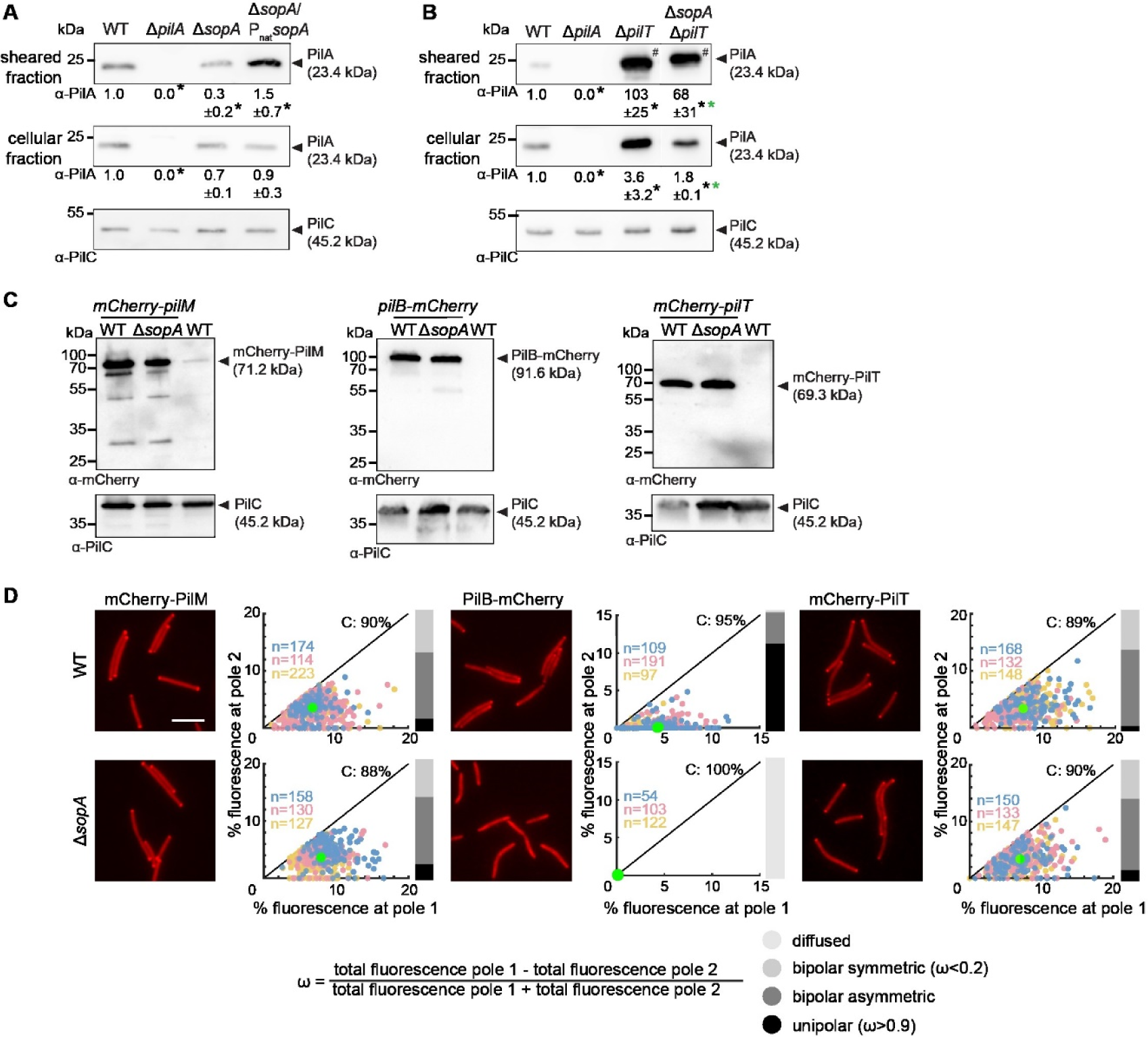
SopA is important for T4aP extension and polar PilB localization. **A.** SopA is important for T4aP formation. T4aP sheared off from 5 mg cells were separated by SDS-PAGE and probed with α-PilA antibodies (top rows). Middle row, protein from total cell extracts of 10^8^ cells was separated by SDS-PAGE and probed with α-PilA antibodies (middle rows), and after stripping, with α-PilC antibodies as a loading control (bottom rows). Numbers below blots indicate PilA levels as the mean ± SD from three biological replicates relative to WT. *, *P*<0.05, two-tailed Student’s *t-*test for samples with equal variances. **B.** SopA is important for T4aP extension. Experiment was done, presented and analyzed as in A. For better comparison, only 10% of T4aP sheared from the hyper-piliated Δ*pilT* strains (#) were loaded. * (black, green), *P*<0.05 compared to WT and the Δ*pilT* mutant, respectively. Gap between lanes, indicate lanes removed for presentation purposes. **C.** Accumulation of mCherry-PilM, PilB-mCherry and mCherry-PilT in the presence and absence of SopA. Protein from total cell extracts of 10^8^ cells was separated by SDS-PAGE and probed with α-mCherry antibodies (top) and after stripping with α-PilC antibodies as a loading control (bottom). All fusion proteins were synthesized from their native locus. **D.** Quantification of the polar localization of mCherry-PilM, PilB-mCherry and mCherry-PilT in the presence and absence of SopA by fluorescence microscopy. Scale bar, 5 μm. In the scatter plots, the percentage of total fluorescence at pole 2 is plotted against the percentage of total fluorescence at pole 1 for all cells with polar cluster(s). Pole 1 is per definition the pole with the highest fluorescence. Individual data points from three independent experiments are shown in three different colors and with the number of cells per replicate indicated in the corresponding colors. Bright green dot, mean fraction of fluorescence at the poles based on all three experiments and including cells with and without clusters. Numbers in the upper right corners, the mean percentage of total cytoplasmic fluorescence based on all three experiments and including cells with and without clusters. Black lines are symmetry lines. For all cells with a cluster(s), an asymmetry index, ω, was calculated as indicated; based on ω values, localization patterns were binned into three categories as indicated; diffuse localization was determined when no polar signal was detected. Bar diagrams to the right, the percentage of cells with a polar localization pattern and diffuse localization according to the color code.

Reduced T4aP formation can be caused by impaired T4aP extension or by increased T4aP retraction. To distinguish these two scenarios, we constructed a Δ*sopA*Δ*pilT* double mutant, which lacks the PilT retraction ATPase, and determined the piliation level of this strain using the shear-off assay. The non-retracting Δ*pilT* mutant, which assembles a very high level of T4aP (66, 67), as well as the Δ*sopA*Δ*pilT* double mutant had significantly higher levels of PilA than WT in the sheared fraction (Fig. 2B). Importantly, the level of PilA in the sheared fraction of the Δ*sopA*Δ*pilT* mutant was significantly lower than in the Δ*pilT* mutant (Fig. 2B). The Δ*pilT* mutant, in agreement with previous observations (7), and the Δ*sopA*Δ*pilT* double mutant both had an increased level of PilA in the cellular fraction (Fig. 2B). Based on these analyses, we conclude that SopA is important but not essential for T4aP extension. By comparison, SgmX is essential for T4aP extension (7). Of note, the observation that the level of PilA in the sheared fraction in the Δ*sopA*Δ*pilT* double mutant is higher than in the Δ*sopA* mutant provides evidence that the Δ*sopA* mutant is able to retract T4aP.

### SopA stimulates polar localization of the PilB extension ATPase

To address how SopA causes a T4aP extension defect, we asked whether lack of SopA causes a defect in the assembly of the T4aPM. The bipolar assembly of the T4aPM core in *M. xanthus* initiates with the OM secretin PilQ (Fig. S1A) then proceeds in an outside-in pathway culminating with the incorporation of PilM (10, 35, 37). Therefore, we used the bipolar localization of a fully active mCherry-PilM fusion synthesized from the native locus (11) (Fig. S1A) as a proxy for the assembly of the T4aPM core. The fusion protein accumulated at the same level in the WT and the Δ*sopA* mutant (Fig. 2C) and localized similarly in the two strains (Fig. 2D). Also, a fully active mCherry-PilT fusion, which was synthesized from the native locus and accumulated at the same level as native PilT (Fig. S4A), accumulated at the same level in the WT and the Δ*sopA* mutant (Fig. 2C) and localized in the same bipolar asymmetric pattern in the two strains (Fig. 2D). By contrast, the polar localization of a partially active PilB-mCherry fusion, which was synthesized from the native site and accumulated at the same level as native PilB (Fig. S4B), was completely abolished in the absence of SopA (Fig. 2D), while it accumulated independently of SopA (Fig. 2C).

We conclude that SopA is not important for the bipolar assembly of the core T4aPM and the polar localization of PilT; however, SopA is essential for polar localization of the PilB extension ATPase. These observations suggest that the defect in T4aP extension caused by lack of SopA is associated with impaired polar localization of PilB. Importantly, SgmX is also essential for polar localization of PilB but not for polar localization of PilM and PilT (7).

### SopA localizes dynamically to the leading cell pole depending on MglA, SgmX and FrzS

To understand the mechanism of SopA in T4aP extension and PilB localization, we asked whether SopA is polarly localized. To this end, we expressed a fully active mVenus-SopA fusion from the native locus (Fig. S5A-B). Using time-lapse fluorescence microscopy and snap-shot image analyses, we observed that mVenus-SopA localized in a unipolar or bipolar asymmetric pattern in all cells and with a large cluster at the leading pole (Fig. 3A-B). During reversals, the polarity of the large cluster was inverted, and after a reversal, it localized at the new leading pole (Fig. 3A).

**Figure 3.**
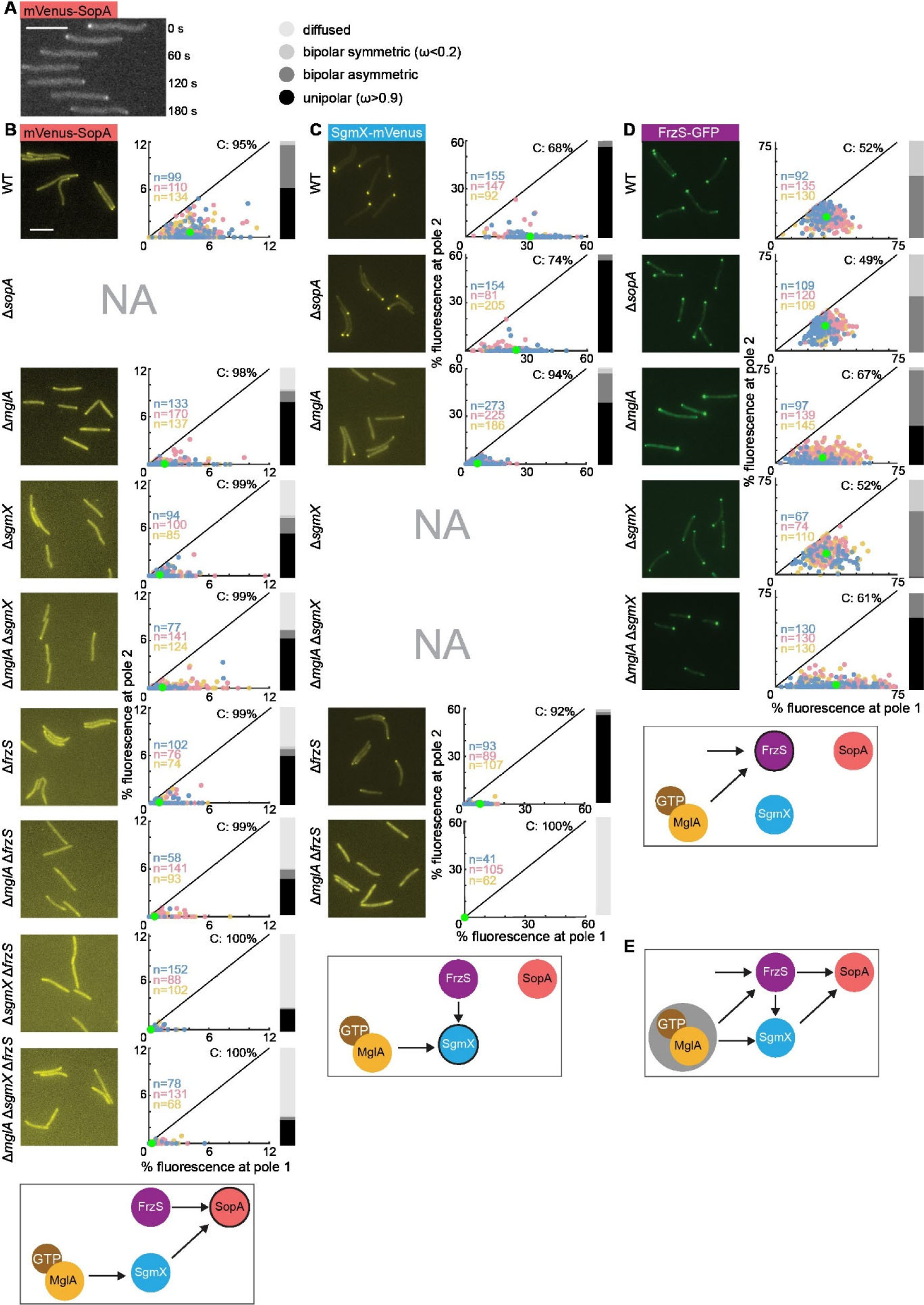
Polar localization of mVenus-SopA, SgmX-mVenus and FrzS-GFP in the presence and absence of MglA, SopA, SgmX and/or FrzS. **A.** mVenus-SopA is dynamically localized with a large cluster at the leading cell pole. Cells were imaged by time-lapse fluorescence microscopy every 30 sec. Scale bar, 5μm. **B-D.** Quantification of the polar localization of mVenus-SopA, SgmX-mVenus and FrzS-GFP. Experiments were done and are presented as in Fig. 2D. All fusion proteins were synthesized from their native locus. Schematics below each row, summarize effects observed. In the schematics, the protein being analyzed for localization is indicated by black circle. **E.** Model of protein interaction network for polar localization of MglA, SgmX, FrzS and SopA. Grey circle surrounding MglA-GTP indicates the polar recruitment of MglA-GTP by the RomR/RomX complex of the polarity module.

Next, we asked whether the polar localization of mVenus-SopA depends on MglA and/or SgmX. In the absence of MglA, fewer cells had polar mVenus-SopA clusters; and in cells with cluster(s), these clusters were of lower intensity than in WT (Fig. 3B). In the absence of SgmX, even fewer cells than in the absence of MglA had polar mVenus-SopA clusters, and in cells with cluster(s), these clusters were of lower intensity than in the absence of MglA (Fig. 3B). Because MglA is important for SgmX polar localization, we determined the localization of mVenus-SopA in a Δ*mglA*Δ*sgmX* double mutant and observed that mVenus-SopA largely localized as in the Δ*sgmX* mutant (Fig. 3B). mVenus-SopA accumulated at the same level in all four strains (Fig. S5B). Altogether, these observations suggest a pathway in which MglA recruits SgmX by direct interaction, and then SgmX, in turn, recruits SopA (Fig. 3B).

In the absence of MglA as well as SgmX, more than 50% of cells still had a polar mVenus-SopA signal. We, therefore, hypothesized that an additional protein would be involved in mVenus-SopA polar recruitment. To test this hypothesis, we took a candidate approach and focused on FrzS, which largely co-occurs with SopA (Fig. S2A). In the Δ*frzS* mutant, fewer cells had polar mVenus-SopA clusters, and in cells with cluster(s), these were of lower intensity than in WT (Fig. 3B). In the Δ*mglA*Δ*frzS* double mutant, most cells did not have a polar cluster and in cells with cluster(s), these were of much lower intensity than in the two strains with a single mutation (Fig. 3B). To test whether SgmX and FrzS have an additive effect on mVenus-SopA polar localization, we generated a Δ*sgmX*Δ*frzS* double mutant. These two mutations had an additive effect on polar mVenus-SopA localization, i.e. most cells did not have polar signal(s) and in the few cells with polar signal(s), these were of very low intensity (Fig. 3B). Finally, in the Δ*mglA*Δ*sgmX*Δ*frzS* triple mutant, mVenus-SopA polar localization was also essentially abolished (Fig. 3B). In all strains, mVenus-SopA accumulated as in WT (Fig. S5B).

We conclude that MglA, SgmX and FrzS are all important for polar localization of mVenus-SopA. The additive effect of the Δ*sgmX* and Δ*frzS* mutations indicate that SgmX and FrzS provide separate inputs to the polar recruitment of mVenus-SopA. Moreover, our data support that in the SgmX pathway, MglA function indirectly to recruit SopA by directly recruiting SgmX, which then recruits SopA (Fig. 3B).

### MglA polar localization is independent of SopA

To address whether SopA is important for MglA polar localization, we imaged the localization of MglA-mVenus in WT and Δ*sopA* cells. MglA-mVenus accumulated (Fig. S6A) and localized similarly in WT and the Δ*sopA* mutant (Fig. S6B). Consistently, SopA was neither important for the accumulation nor the polar localization of RomR and MglB, two key proteins of the polarity module (Fig. S1B; Fig. S6A-B). We conclude that SopA acts downstream of the polarity module to stimulate polar localization of PilB and, thereby, T4aP extension and T4aP-dependent motility.

### SgmX polar localization depends on MglA and FrzS but not on SopA

To further understand the interplay between MglA, SgmX, FrzS and SopA for polar localization, we explored the localization of SgmX. In agreement with previous observations (7, 46), a fully active SgmX-mVenus fusion localized in a highly unipolar pattern in WT and this polar localization was strongly reduced in the Δ*mglA* mutant (Fig. 3C); however, it was not affected in the Δ*sopA* mutant (Fig. 3C). In the absence of FrzS, SgmX-mVenus polar localization was also strongly reduced (Fig. 3C) in agreement with recent observations (58). Moreover, in the Δ*mglA*Δ*frzS* double mutant, SgmX-mVenus polar localization was completely abolished. In all strains, SgmX-mVenus accumulated as in WT (Fig. S5C). These observations suggest that SgmX polar recruitment depends on two pathways, one involves MglA and one involves FrzS (Fig. 3C).

### FrzS polar localization depends on MglA but not on SgmX and SopA

Next, we explored polar FrzS localization. To this end, we used a fully active FrzS-GFP fusion synthesized from the native locus [(68); Fig. S5D]. In agreement with previous observations (53, 55, 68), FrzS-GFP localized in a bipolar asymmetric pattern in WT, and this localization was reduced and shifted to more asymmetric in the absence of MglA (Fig. 3D). FrzS-GFP localization was not affected by the lack of SopA (Fig. 3D). Similarly, FrzS-GFP was not affected by the lack of SgmX (Fig. 3D). Finally, in the Δ*mglA*Δ*sgmX* double mutant, FrzS-GFP localized in the more asymmetric pattern observed in the Δ*mglA* mutant (Fig. 3D). We conclude that MglA is important for polar FrzS-GFP localization while SgmX and SopA are not (Fig. 3D). We also note that in the Δ*mglA*Δ*sgmX* double mutant, FrzS-GFP formed polar clusters in all cells, documenting that MglA is not the only polar recruitment factor of FrzS. In all tested strains, FrzS-GFP accumulated as in WT (Fig. S5E).

### A highly interconnected protein interaction network establishes the polar localization of MglA, SgmX, FrzS and SopA

Collectively, the MglA-mVenus, mVenus-SopA, SgmX-mVenus and FrzS-GFP localization patterns described in Fig. 3B-D and Fig. S6B together with previous findings suggest that multiple interactions between these four proteins establish a highly interconnected network that results in their polar localization (Fig. 3E). In this network, two proteins can localize polarly in the absence of the three other proteins, thereby establishing the basis for the recruitment of the remaining two proteins. The first protein is MglA, and neither FrzS (53) nor SgmX (7) nor SopA (Fig. S6B) are important for MglA polar localization, suggesting that MglA is only recruited to the pole *via* the RomR/RomX complex of the polarity module (Fig. 3E). The second protein is FrzS, which can localize polarly independently of MglA, SgmX and SopA (Fig. 3D). MglA further stimulates FrzS polar localization (Fig. 3D). Downstream of MglA and FrzS, these two proteins can separately recruit SgmX, i.e. MglA can recruit SgmX in the absence of FrzS and *vice versa* (Fig. 3C). Finally, FrzS and SgmX can separately recruit SopA (Fig. 3B). Conversely, SopA neither affects MglA, SgmX nor FrzS polar localization. In this pathway, the effect of MglA on SopA localization is indirect and depends on the effect of MglA on FrzS and SgmX polar recruitment.

### FrzS is important for T4aP extension and polar PilB localization

To examine how the protein interaction network for polar localization of MglA, SgmX, FrzS and SopA relates to T4aP formation and T4aP-dependent motility, we first characterized T4aP-dependent motility, T4aP formation and PilB localization in the Δ*frzS* mutant. In agreement with previous observations (57), the Δ*frzS* mutant had significantly reduced T4aP-dependent motility (Fig. 4A). Moreover, and in agreement with FrzS being important for SgmX and SopA polar localization (Fig. 3E), the Δ*frzS* mutant had significantly reduced PilA in the sheared T4aP fraction (Fig. 4B), and the Δ*frzS*Δ*pilT* double mutant had reduced PilA in the sheared T4aP fraction compared to the Δ*pilT* mutant (Fig. 4C). Furthermore, PilB polar localization was strongly reduced but not abolished in the Δ*frzS* mutant (Fig. 4D). These observations suggest that the defect in T4aP extension caused by lack of FrzS is caused by the reduced polar localization of PilB. Because FrzS is important for polar localization of SgmX and SopA (Fig. 3E), which are, in turn, essential for PilB polar localization, these observations support that the effect of lack of FrzS on T4aP-dependent motility, T4aP formation and polar localization of PilB are mediated via its effect on SgmX and SopA polar localization.

**Figure 4.**
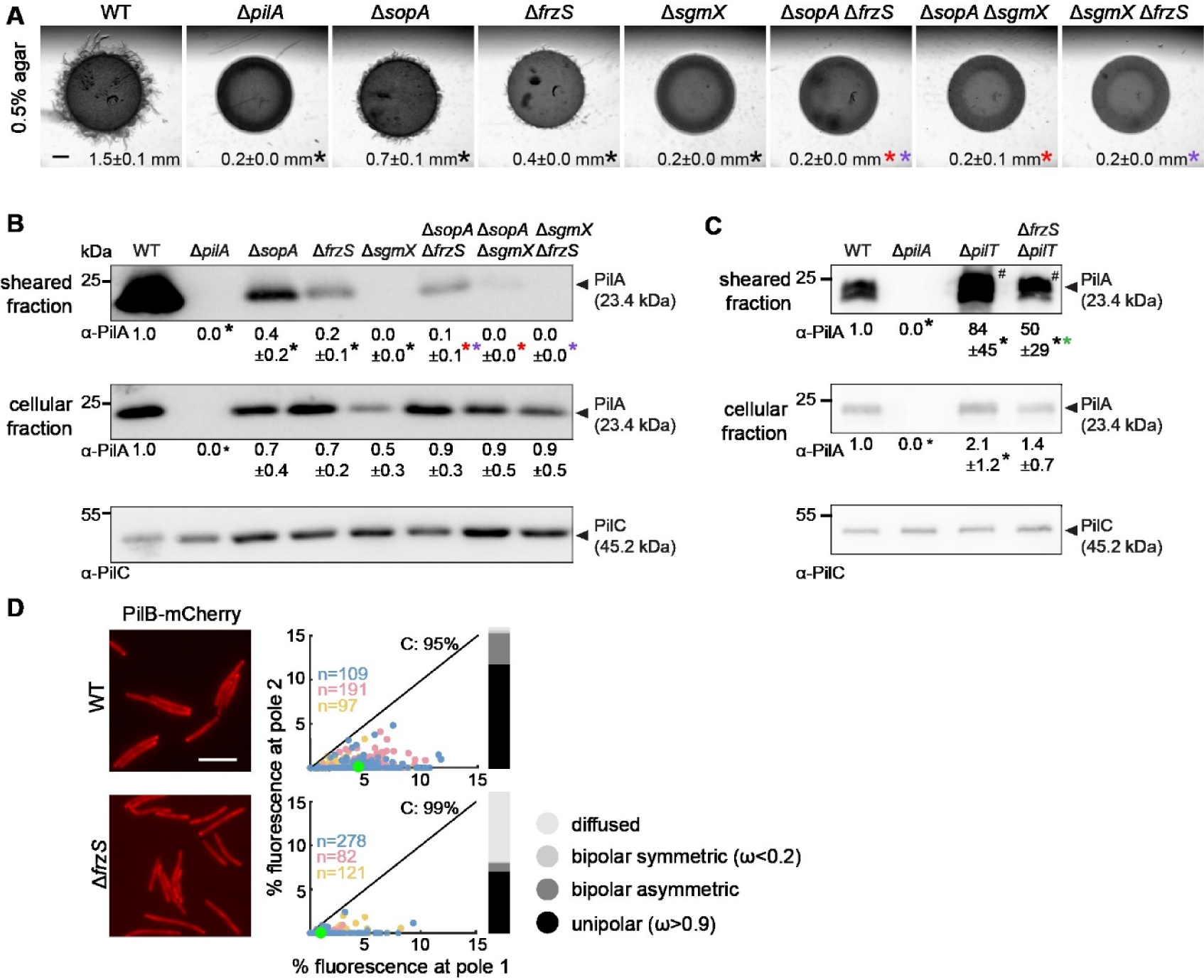
Combinatorial effect of SgmX, FrzS and SopA on T4aP-dependent motility and T4aP formation. **A.** Effect of SgmX, FrzS and/or SopA on T4aP-dependent motility. Cells were incubated on 0.5% agar supplemented with 0.5% CTT. Scale bar, 1mm. Numbers, colony expansion in mm in 24h as mean ± SD from three biological replicates; * (black, red, purple) *P*<0.05, two-tailed Student’s *t-*test for samples with equal variances compared to WT, the Δ*sopA* mutant and the Δ*frzS* mutant, respectively. **B-C.** Effect of SgmX, FrzS and/or SopA on T4aP formation. Experiments were done and data presented as in Fig. 2A-B, excepti that in B T4aP sheared off from 7.5 mg cells were loaded. * (black, red, purple, green), *P*<0.05, two-tailed Student’s *t-*test for samples with equal variances compared to WT, the Δ*sopA* mutant, the Δ*frzS* mutant and the Δ*pilT* mutant, respectively. **D.** FrzS is important for polar localization of PilB-mCherry. Experiment was done and data presented as in Fig. 2D.

### The MglA/SgmX/FrzS/SopA interaction network establishes different levels of T4aP-formation and T4aP-dependent motility

Having demonstrated that the polar localization of SgmX, FrzS and SopA is governed by an intricate set of interactions, we hypothesized that these three proteins would have differential effects on T4aP formation and, thus, T4aP-dependent motility. To test this hypothesis, we compared the defects in T4aP formation and T4aP-dependent motility in the Δ*sgmX*, Δ*frzS* and Δ*sopA* mutants and the three double mutants. This comparison revealed that the amount of PilA in the sheared fraction in the six mutants followed a gradient, i.e. the Δ*sopA* mutant had significantly reduced PilA in the sheared fraction, the Δ*frzS* mutant was even more reduced, the Δ*sopA*Δ*frzS* mutant was even more strongly reduced, and PilA in the sheared fraction was undetectable in the Δ*sgmX* mutant and in the two Δ*sopA*Δ*sgmX* and Δ*frzS*Δ*sgmX* double mutants (Fig. 4B). Notably, with the exception of the Δ*sopA*Δ*frzS* mutant, the defects in T4aP formation correlated with the level of T4aP-dependent motility in the different mutants, i.e. it was significantly reduced in the Δ*sopA* mutant, even more strongly reduced in the Δ*frzS* mutant, and abolished in the Δ*sgmX* mutant an the three double mutants (Fig. 4A). The Δ*sopA*Δ*frzS* mutant, which was ∼10-fold reduced in the amount of PilA in the sheared fraction compared to WT, did not detectably display T4aP-dependent motility under these test conditions.

### SopA interacts directly with SgmX

SgmX directly interacts with MglA (7, 46) and FrzS (58). Moreover, it has been suggested that MglA interacts directly with FrzS based on *in vivo* pull-down experiments (55). To shed light on whether SopA interacts directly with SgmX and/or FrzS, we used bacterial adenylate cyclase-based two hybrid (BACTH) analyses (69) with full-length SopA, SgmX and FrzS proteins. We observed that SgmX as well as FrzS self-interacted (Fig. 5; Fig. S7) in agreement with the observations that purified SgmX and FrzS are both likely dimeric (7, 57). Moreover, we observed interactions between SgmX and FrzS as well as between SgmX and SopA but not between SopA and FrzS (Fig. 5; Fig. S7).

**Figure 5.**
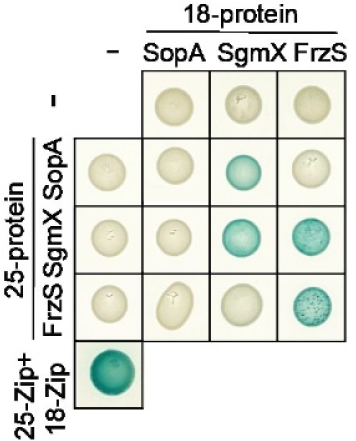
BACTH assay for SgmX, FrzS and SopA interactions. Full-length SgmX, FrzS and SopA were fused to the C-terminus of T25 and T18. Lower left corner, T25-Zip + T18-Zip positive control.

## Discussion

In this study, we addressed how T4aP formation is regulated in the rod-shaped cells of *M. xanthus*. Altogether, the detailed quantification of protein localization and T4aP formation supports a model in which the four proteins MglA, SgmX, FrzS and SopA establish a highly interconnected protein interaction network to regulate T4aP formation (Fig. 6). In this network, the small GTPase MglA is recruited to the leading pole *via* the RomR/RomX complex of the polarity module. MglA and its downstream effector protein SgmX are required and sufficient for the unipolar formation of T4aP and jointly bring about a low level of T4aP formation. By contrast, FrzS and SopA are dispensable for T4aP formation, and these two proteins function to stimulate the MglA/SgmX pathway for T4aP formation. In agreement with previous observations, FrzS is recruited to the leading pole by MglA-dependent and MglA-independent mechanisms. At this pole, FrzS stimulates SgmX polar localization and, thus, T4aP formation. In the case of SopA, it is separately recruited to the leading pole by SgmX and FrzS, where it stimulates the MglA/SgmX pathway for T4aP formation. Because SgmX and SopA are essential for the polar localization of the PilB extension ATPase while FrzS is important, we propose that the output of this pathway is to stimulate PilB interaction with the cytoplasmic base of the core T4aPM (Fig. 6), thereby licensing T4aP formation. Because SopA does not affect the polar localization of MglA, SgmX and FrzS, we suggest that SopA stimulates the function of SgmX in PilB polar recruitment (Fig. 6).

**Figure 6.**
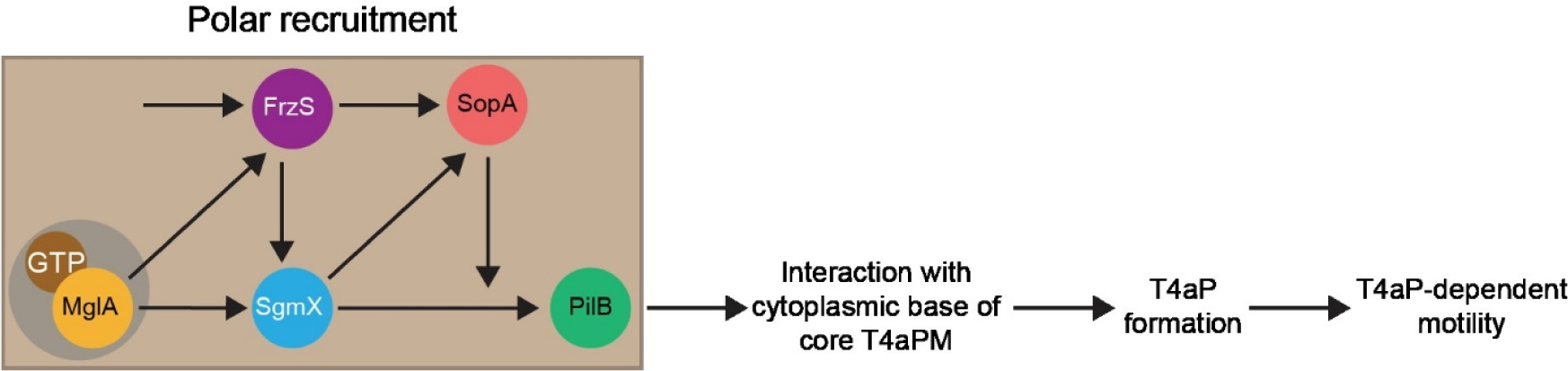
Model of protein interaction network for combinatorial regulation of T4aP formation and T4aP-dependent motility in *M. xanthus*. Light brown box indicates interactions that stimulate polar recruitment of proteins; grey circle surrounding MglA-GTP indicates the polar recruitment of MglA-GTP by the RomR/RomX complex of the polarity module.

The detailed quantification of T4aP formation in different mutants provides evidence that the MglA/SgmX/FrzS/SopA interaction network allows for combinatorial regulation of the level of T4aP formation. Specifically, this network can distinguish at least five input states that generate five corresponding output states with five discrete levels of T4aP formation: (i) in the absence of MglA and SgmX, no T4aP are formed (7), (ii) in the presence of only MglA and SgmX, a low level of T4aP is assembled, (iii) in the presence of MglA, SgmX and FrzS, the level is increased, (iv) in the presence of MglA, SgmX and SopA, an even higher level of T4aP is assembled, and, finally, (v) in the presence of all four proteins, the WT level of T4aP formation is accomplished. Thus, this pathway allows the regulation of the number of T4aP by integrating the input from MglA, FrzS and SopA on the central protein SgmX. Under the conditions of the assay for T4aP-dependent motility, the defects in T4aP formation correlated with the level of T4aP-dependent motility in the different mutants except for the Δ*sopA*Δ*frzS* mutant. This mutant had a ∼10-fold reduced amount of PilA in the sheared fraction compared to WT and did not display T4aP-dependent motility, suggesting that the number of T4aP in this mutant is too low to enable the pulling of cells across the surface used in the assay for T4aP-dependent motility.

SgmX with its 14 TPRs contains three functional regions (7, 46, 58). The eight N-terminal TPRs mediate the activation of T4aP-dependent motility, the three middle TPRs engage in the interaction to FrzS, and the three C-terminal TPRs in the interaction to MglA (46, 58). FrzS is a pseudo-response regulator with an N-terminal receiver domain, which lacks critical residues for phosphorylation, and a large C-terminal coiled-coil domain (56, 57). The pseudo-receiver domain of FrzS interacts with SgmX (58) while the C-terminal coiled coil is sufficient for polar localization of FrzS (70). Previously, MglA was suggested to interact directly with FrzS (55); however, it is not known how MglA might interact with FrzS. Using a BACTH assay, we observed that SopA interacts directly with SgmX, however, we did not detect an interaction between SopA and FrzS. Based on the dissection of SgmX by Bautista et al. and Mercier et al. (46, 58), we suggest that the eight N-terminal TPRs of SgmX are involved in the polar recruitment of PilB to the T4aPM. PilB interacts directly with PilM and PilC at the cytoplasmic base of the T4aPM [(Fig. S1; (15, 71, 72)]. However, direct interactions between SgmX and PilB and/or PilM have not been detected (7). Therefore, important goals for the future will be to determine how SgmX stimulates the interaction of PilB with the cytoplasmic base of the T4aPM and how SopA might stimulate this interaction. Interestingly, despite PilB not being polarly localized in the absence of SopA, the Δ*sopA* mutant still makes T4aP, suggesting that the formation of a visible polar PilB cluster may not fully reflect the interaction of PilB with the cytoplasmic base of the core T4aPM.

In other bacteria the regulation of T4aP formation also centers on the PilB extension ATPase. Specifically, in *Vibrio cholerae* and *Clostridium perfringens,* the second messenger c-di-GMP binds directly to the MshE and PilB2 ATPase, respectively to stimulate T4aP formation (73–75). In *Xanthomonas axonopodis* pv. citri, c-di-GMP binds to the effector protein FimX, which then interacts with PilZ that, in turn, interacts with PilB, likely to stimulate T4aP formation (23, 76, 77). Similarly, in *Pseudomonas aeruginosa*, the c-di-GMP binding effector proteins FimX stimulate T4aP formation by interacting directly with PilB (25).

The genetic and cell biological analyses demonstrate that the MglA/SgmX/FrzS/SopA network for T4aP formation is able to distinguish different input states with the formation of discrete levels of T4aP. However, the pathway is based on complete loss of function of MglA, SgmX, FrzS and SopA. Therefore, in the future, it will be interesting to investigate under which physiologically conditions these four proteins have altered accumulation and/or localization. In this context, we note that biosynthetic mutants unable to synthesize the secreted polysaccharide exopolysaccharide (EPS) have reduced but not abolished T4aP formation (78). This defect is caused by reduced T4aP extension and not increased retraction (78), but it is not known what causes this extension defect. MglA, SgmX, FrzS and SopA accumulate at WT levels in an Δ*epsZ* mutant (79) that lacks the phosphoglycosyl transferase EpsZ that initiates EPS biosynthesis (78). In the future, it will be of interest to determine the localization of MglA, SgmX, FrzS and SopA in EPS biosynthetic mutants.

## Acknowledgement

We thank María Pérez-Burgos and Marco Herfurth for many helpful discussions. This work was supported by the Max Planck Society.

## Availability of data and materials

The authors declare that all data supporting this study are available within the article and its Supplementary Information files. All materials used in the study are available from the corresponding author.

## Materials & Methods

### Cell growth and construction of strains

Strains, plasmids and primers used in this work are listed in Table 1, 2 and S2, respectively. All *M. xanthus* strains are derivatives of the DK1622 WT strain (38). *M. xanthus* was grown at 32°C in 1% casitone broth (CTT) (80) or on 1.5% agar supplemented with 1% CTT and kanamycin (50µg mL^-1^) or oxytetracycline (10µg mL^-1^) as appropriate. In-frame deletions were generated as described (81). Plasmids were introduced in *M. xanthus* by electroporation and integrated by homologous recombination at the native locus or by site-specific recombination at the Mx8 *attB* site. All in-frame deletions and plasmid integrations were verified by PCR. Plasmids were propagated in *Escherichia coli* TOP10 (F^-^, *mcrA*, Δ(*mrr-hsd*RMS-*mcr*BC), φ80*lac*ZΔM15, Δ*lac*X74, *deoR, recA1, araD139*, Δ(*ara-leu*)7679, *galU, galK, rpsL, endA1, nu*pG). *E. coli* was grown in Lysogeny broth (LB) or on plates containing LB supplemented with 1.5% agar at 37°C with added antibiotics when appropriate (82). All DNA fragments generated by PCR were verified by sequencing.

**Table 1.** *M. xanthus* strains used in this work.

| Strain | Genotype | Reference |
| --- | --- | --- |
| DK1622 | Wild-type | (38) |
| DK10410 | $\Delta pilA$ | (92) |
| SA5293 | $\Delta aglQ$ | (93) |
| SA9828 | $\Delta sopA$ | This work |
| SA9829 | $\Delta sopA \Delta pilA$ | This work |
| SA9830 | $\Delta sopA \Delta aglQ$ | This work |
| SA9835 | $\Delta sopA$ P<sub>nat</sub>*sopA* (*attB*::pMO28) | This work |
| DK10409 | $\Delta pilT$ | (67) |
| SA9859 | $\Delta sopA \Delta pilT$ | This work |
| SA7896 | *mCherry-pilM* | (11) |
| SA9300 | *pilB-mCherry* | This work |
| SA9307 | *mCherry-pilT* | This work |
| SA9837 | *mCherry-pilM* $\Delta sopA$ | This work |
| SA9853 | *pilB-mCherry* $\Delta sopA$ | This work |
| SA9854 | *mCherry-pilT* $\Delta sopA$ | This work |
| SA8185 | *mgIA-mVenus* | (49) |
| SA3963 | *mgIB-mCherry* | (54) |
| SA7507 | *romR-mCherry* | (49) |
| SA9845 | *mgIA-mVenus* $\Delta sopA$ | This work |
| SA9842 | *mgIB-mCherry* $\Delta sopA$ | This work |
| SA9846 | *romR-mCherry* $\Delta sopA$ | This work |
| SA9848 | *mVenus-sopA* | This work |
| SA9852 | *mVenus-sopA* $\Delta mgIA$ | This work |
| SA9855 | *mVenus-sopA* $\Delta sgmX$ | This work |
| SA9857 | *mVenus-sopA* $\Delta frzS$ | This work |
| SA9867 | *mVenus-sopA* $\Delta frzS \Delta mgIA$ | This work |
| SA9868 | *mVenus-sopA* $\Delta sgmX \Delta mgIA$ | This work |
| SA9861 | *mVenus-sopA* $\Delta sgmX \Delta frzS$ | This work |
| SA9869 | *mVenus-sopA* $\Delta sgmX \Delta frzS \Delta mgIA$ | This work |
| SA7164 | $\Delta sgmX$ | (7) |
| SA7195 | *sgmX-mVenus* | (7) |
| SA9851 | *sgmX-mVenus* $\Delta sopA$ | This work |
| SA7196 | *sgmX-mVenus* $\Delta mgIA$ | (7) |
| SA9885 | *sgmX-mVenus* $\Delta frzS$ | This work |
| SA9886 | *sgmX-mVenus* $\Delta mgIA \Delta frzS$ | This work |
| SA9318 | $\Delta frzS$ | This work |
| SA9877 | $\Delta pilT \Delta frzS$ | This work |
| SA9870 | *pilB-mCherry* $\Delta frzS$ | This work |
| SA9879 | *frzS-gfp* | This work |
| SA9880 | *frzS-gfp* $\Delta sopA$ | This work |
| SA9881 | *frzS-gfp* $\Delta mgIA$ | This work |
| SA9882 | *frzS-gfp* $\Delta sgmX$ | This work |
| SA9883 | *frzS-gfp* $\Delta sgmX \Delta mgIA$ | This work |
| SA9860 | $\Delta sopA \Delta frzS$ | This work |
| SA9856 | $\Delta sopA \Delta sgmX$ | This work |

**Table 2.**
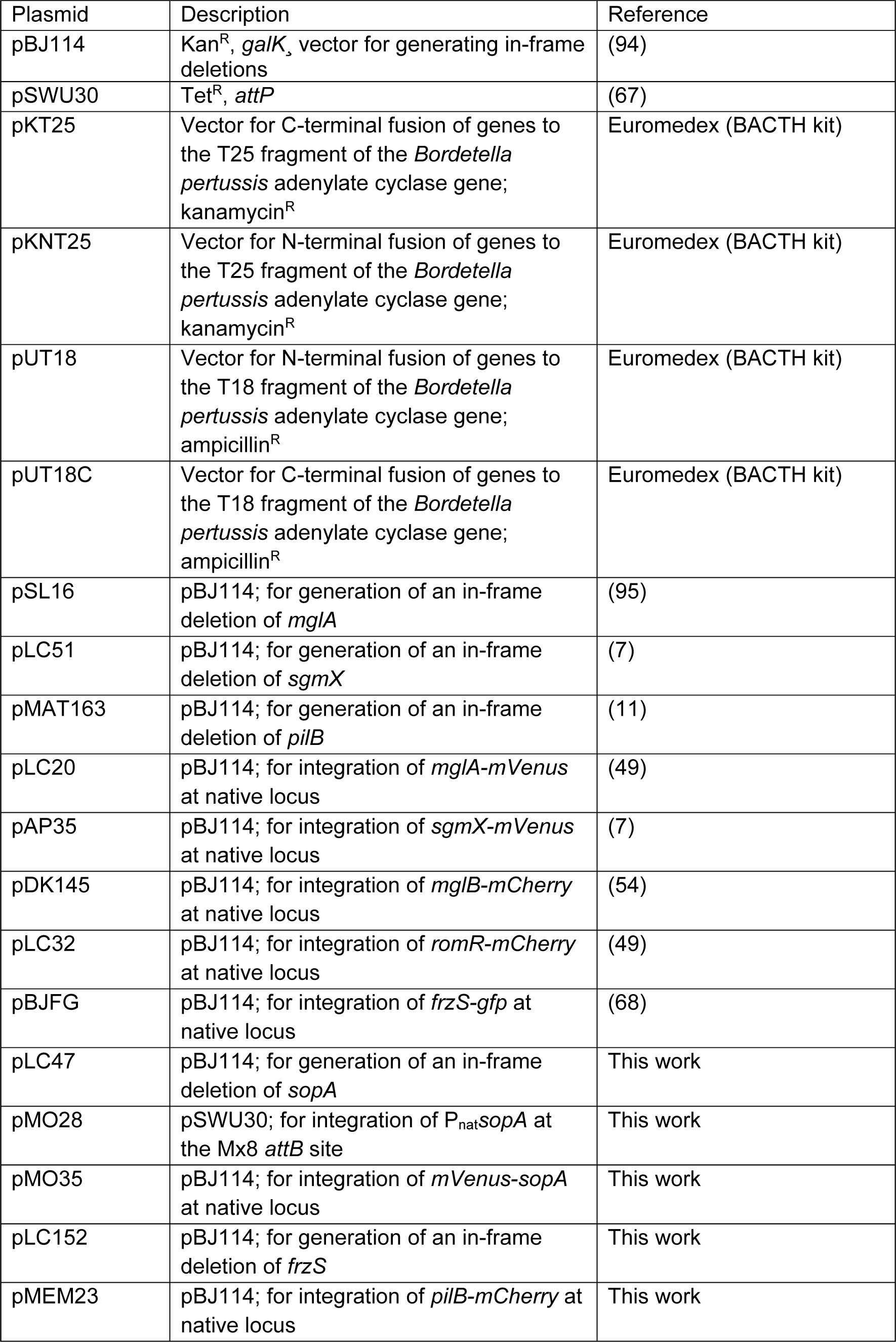
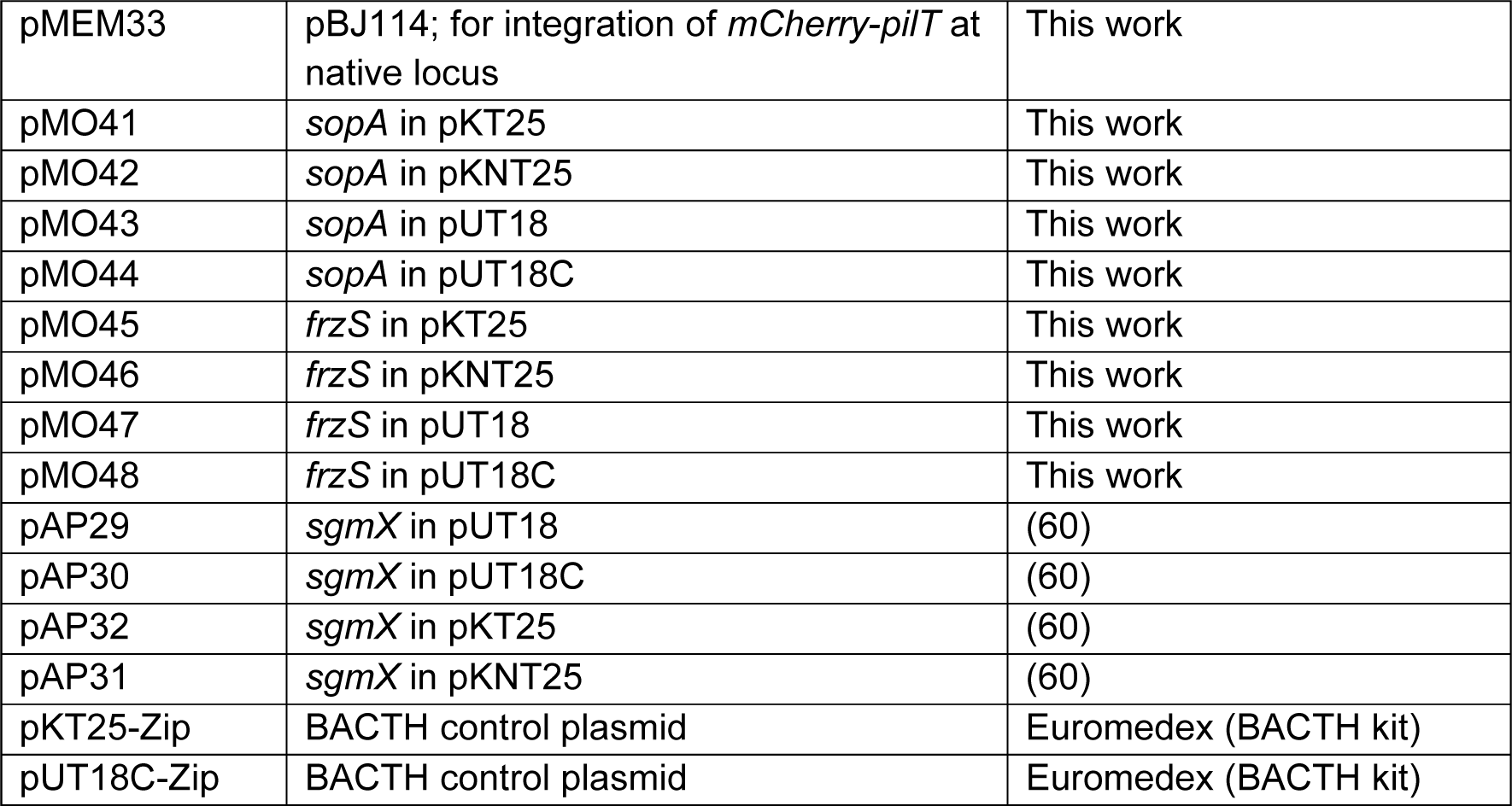
Plasmids used in this work.

| Plasmid | Description | Reference |
| --- | --- | --- |
| pBJ114 | Kan<sup>R</sup>, *galK*, vector for generating in-frame deletions | (94) |
| pSWU30 | Tet<sup>R</sup>, *attP* | (67) |
| pKT25 | Vector for C-terminal fusion of genes to the T25 fragment of the *Bordetella pertussis* adenylate cyclase gene; kanamycin<sup>R</sup> | Euromedex (BACTH kit) |
| pKNT25 | Vector for N-terminal fusion of genes to the T25 fragment of the *Bordetella pertussis* adenylate cyclase gene; kanamycin<sup>R</sup> | Euromedex (BACTH kit) |
| pUT18 | Vector for N-terminal fusion of genes to the T18 fragment of the *Bordetella pertussis* adenylate cyclase gene; ampicillin<sup>R</sup> | Euromedex (BACTH kit) |
| pUT18C | Vector for C-terminal fusion of genes to the T18 fragment of the *Bordetella pertussis* adenylate cyclase gene; ampicillin<sup>R</sup> | Euromedex (BACTH kit) |
| pSL16 | pBJ114; for generation of an in-frame deletion of *mglA* | (95) |
| pLC51 | pBJ114; for generation of an in-frame deletion of *sgmX* | (7) |
| pMAT163 | pBJ114; for generation of an in-frame deletion of *pilB* | (11) |
| pLC20 | pBJ114; for integration of *mglA-mVenus* at native locus | (49) |
| pAP35 | pBJ114; for integration of *sgmX-mVenus* at native locus | (7) |
| pDK145 | pBJ114; for integration of *mglB-mCherry* at native locus | (54) |
| pLC32 | pBJ114; for integration of *romR-mCherry* at native locus | (49) |
| pBJFG | pBJ114; for integration of *frzS-gfp* at native locus | (68) |
| pLC47 | pBJ114; for generation of an in-frame deletion of *sopA* | This work |
| pMO28 | pSWU30; for integration of P<sub>nat</sub>*sopA* at the Mx8 *attB* site | This work |
| pMO35 | pBJ114; for integration of *mVenus-sopA* at native locus | This work |
| pLC152 | pBJ114; for generation of an in-frame deletion of *frzS* | This work |
| pMEM23 | pBJ114; for integration of *pilB-mCherry* at native locus | This work |
| pMEM33 | pBJ114; for integration of *mCherry-pilT* at native locus | This work |
| pMO41 | *sopA* in pKT25 | This work |
| pMO42 | *sopA* in pKNT25 | This work |
| pMO43 | *sopA* in pUT18 | This work |
| pMO44 | *sopA* in pUT18C | This work |
| pMO45 | *frzS* in pKT25 | This work |
| pMO46 | *frzS* in pKNT25 | This work |
| pMO47 | *frzS* in pUT18 | This work |
| pMO48 | *frzS* in pUT18C | This work |
| pAP29 | *sgmX* in pUT18 | (60) |
| pAP30 | *sgmX* in pUT18C | (60) |
| pAP32 | *sgmX* in pKT25 | (60) |
| pAP31 | *sgmX* in pKNT25 | (60) |
| pKT25-Zip | BACTH control plasmid | Euromedex (BACTH kit) |
| pUT18C-Zip | BACTH control plasmid | Euromedex (BACTH kit) |

### Motility assays and determination of reversal frequency

Population-based motility assays were done as described (62). Briefly, *M. xanthus* cells from exponentially growing cultures were harvested at 4000 *g* for 10 min at room temperature (RT) and resuspended in 1% CTT to a calculated density of 7×10^9^ cells mL^-1^. 5µL aliquots of cell suspensions were placed on 0.5% agar plates supplemented with 0.5% CTT for T4aP-dependent motility and 1.5% agar plates supplemented with 0.5% CTT for gliding motility and incubated at 32°C. At 24 h, colony edges were visualized using a Leica M205FA stereomicroscope and imaged using a Hamamatsu ORCA-flash V2 Digital CMOS camera (Hamamatsu Photonics) using the LASX software (Leica Microsystems). For higher magnifications of cells at colony edges on 1.5% agar, cells were visualized using a Leica DMi8 inverted microscope and imaged with a Leica DFC9000 GT camera. Single cells were tracked as described (49). Briefly, for T4aP-dependent motility, 5µL of exponentially growing cultures were placed in a 24-well polystyrene plate (Falcon). After 10 min at RT, cells were covered with 200 µL 1% methylcellulose in MMC buffer (10mM MOPS (3-(*N*-morpholino) propanesulfonic acid) pH 7.6, 4mM MgSO_4_, 2mM CaCl_2_), and incubated at RT for 30 min. Subsequently, cells were visualized for 15 min at 20 sec intervals at RT using a Leica DMi8 inverted microscope with a Leica DFC9000 GT camera and using the LASX software (Leica Microsystems). Individual cells were tracked using Metamorph 7.5 (Molecular Devices) and ImageJ 1.52b (83) and then the speed of individual cells per 20 sec interval as well as the number of reversals per cell per 15 min calculated. For gliding, 3µL of exponentially growing cultures were placed on 1.5% agarose plates supplemented with 0.5% CTT, covered by a cover slide and incubated at 32°C. After 4 to 6h, cells were observed for 15 min at 30 sec intervals at RT as described, speed per 30 sec interval as well as the number of reversals per 15 min calculated. In both assays, only cells that moved for the entire recording period were included.

### Immunoblot analysis

Immunoblot analysis was done as described (82). Rabbit polyclonal α-PilA (11) (dilution 1:3000), α-PilC (34) (dilution 1:5000), α-mCherry (Biovision, dilution 1:15000), α-PilT (66) (dilution 1:2000) and α-PilB (66) were used together with horseradish peroxidase-conjugated goat anti-rabbit immunoglobulin G (Sigma) as a secondary antibody (dilution 1:10000). Monoclonal mouse antibodies were used to detect GFP-tagged proteins (Roche) (dilution 1:2000) together with horseradish peroxidase conjugated sheep anti-mouse immunoglobulin G (GE Healthcare) as a secondary antibody (dilution 1:2000). Blots were developed using Luminata Crescendo Western HRP substrate (Millipore) and visualized using a LAS-4000 luminescent image analyzer (Fujifilm). Proteins were separated by SDS-PAGE as described (82).

### T4aP shearing assays

Pili were sheared of *M. xanthus* cells using a protocol based on the procedure of (67). Briefly, cells grown on 1% CTT, 1.5% agar plates for 2-3 days were gently scraped off the agar and resuspended in pili resuspension buffer (100 mM Tris-HCl pH 7.6, 150 mM NaCl) (1 mL per 60 mg cells). Cell suspensions were vortexed for 10 min at highest speed. Cells from a 100 µL aliquot were harvested, the pellet dissolved in 100 µL SDS lysis buffer (10% (v/v) glycerol, 50 mM Tris-HCl pH 6.8, 2 mM EDTA, 2% (w/v) SDS, 100 mM DTT, 0.01% bromphenol blue) and immediately denatured at 95°C for 5 min. The remaining suspension was centrifuged for 20 min at 13,000 *g* at 4°C. The supernatant removed and centrifuged twice for 10 min at 13,000 *g* at 4°C to remove cell debris. T4aP in the cell-free supernatant were precipitated by adding 10× pili precipitation buffer (final concentrations: 100 mM MgCl_2_, 500 mM NaCl, 2% PEG 6000) for at least 2 h at 4°C. The solution was centrifuged for 30 min at 13,000 *g* at 4°C and the pellet suspended in SDS lysis buffer (1 µL per mg vortexed cells). T4aP sheared and purified from the same amount of cells were loaded and separated by SDS-PAGE.

### Bacterial Adenylate Cyclase-Based Two-Hybrid (BACTH) assays

BACTH assays were performed according to the manufacturer’s protocol (Euromedex). Briefly, plasmids encoding full-length SgmX, FrzS or SopA fused N-terminally or C-terminally to the T25 or T18 *Bordetella pertussis* adenylate cyclase (CyaA) fragments were transformed into *E. coli* BTH101 (F-*cya*-99 *araD*139 *galE*15 *galK*16 *rpsL1* (Str^r^) *hsdR*2 *mcrA*1 *mcrB*1) alone or in pairs. As a positive control, BTH101 co-transformed with the plasmids pKT25-zip and pUT18C-zip were used. Transformed cells were incubated at 30°C for 24 h. cAMP production by reconstituted CyaA was qualitatively assessed by the formation of blue color as a read out for protein-protein interactions on LB agar supplemented with 40 µg ml^−1^ 5-bromo-4-chloro-3-indolyl-β-d-galactopyranoside and 0.5 mM isopropyl-β-D-thiogalactopyranosid (IPTG).

### Fluorescence microscopy and image analysis

For fluorescence microscopy, exponentially growing cells were placed on slides containing a thin pad of 1% SeaKem LE agarose (Cambrex) with TPM buffer (10mM Tris-HCl pH 7.6, 1mM KH_2_PO_4_ pH 7.6, 8mM MgSO_4_) and 0.2% CTT, and covered with a coverslip. After 30 min at 32°C, cells were visualized using a temperature-controlled Leica DMi8 inverted microscope and phase contrast and fluorescence images acquired using a Hamamatsu ORCA-flash V2 Digital CMOS camera and the LASX software (Leica Microsystems). For time-lapse recordings, cells were imaged for 15 min using the same conditions. Microscope images were processed with Fiji (84) and cell masks determined using Oufti (85) and manually corrected when necessary. To precisely quantify the localization of fluorescently-labelled proteins, we used Matlab R2020a (The MathWorks) in an established analysis pipeline (51) in which the output for each cell is total cellular fluorescence and fluorescence in clusters at each pole. Briefly, cells were segmented, and polar clusters were identified as having an average fluorescence signal of 2 SD above the mean cytoplasmic fluorescence and a size of three or more pixels. Pole 1 was assigned to the pole with the highest fluorescence. For each cell with polar clusters, an asymmetry index (ω) was calculated as:

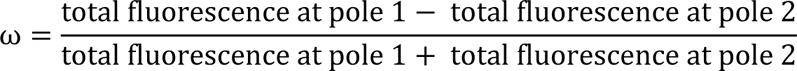

The localization patterns were binned from the ω values as follows: unipolar (ω>0.9), bipolar asymmetric (0.9≥ω≥0.2) and bipolar symmetric (ω<0.2). Diffuse localization was determined when no polar signal was detected. Data points for individual cells were plotted in scatterplots. For calculating mean fraction of polar and cytoplasmic fluorescence, cells with and without clusters were included.

### Bioinformatics

The search of the STRING database (59) for proteins that co-occur with SgmX was conducted October 2016. Sequence alignments were generated using ClustalOmega (86) with default parameters and alignments were visualized with Jalview (87). Protein domains were identified using Interpro (88). Orthologs were identified using the KEGG SSDB database (89). % similarity/identity between proteins were calculated using EMBOSS Needle software (pairwise sequence alignment) (90). Phylogenetic trees were prepared in MEGA7 (91) using the Neighbor-Joining method.

### Statistics

Statistics were performed using a two-tailed Student’s *t*-test for samples with equal variances.

## References

1. Wadhwa N, Berg HC. 2022. Bacterial motility: machinery and mechanisms. Nat Rev Microbiol 20:161–173.

2. Craig L, Forest KT, Maier B. 2019. Type IV pili: dynamics, biophysics and functional consequences. Nat Rev Microbiol 17:429–440.

3. Merz AJ, So M, Sheetz MP. 2000. Pilus retraction powers bacterial twitching motility. Nature 407:98–102.

4. Skerker JM, Berg HC. 2001. Direct observation of extension and retraction of type IV pili. Proc Natl Acad Sci U S A 98:6901–4.

5. Maier B, Potter L, So M, Long CD, Seifert HS, Sheetz MP. 2002. Single pilus motor forces exceed 100 pN. Proc Natl Acad Sci USA 99:16012–16017.

6. Clausen M, Jakovljevic V, Søgaard-Andersen L, Maier B. 2009. High force generation is a conserved property of type IV pilus systems. J Bacteriol 191:4633–4638.

7. Potapova A, Carreira LAM, Søgaard-Andersen L. 2020. The small GTPase MglA together with the TPR domain protein SgmX stimulates type IV pili formation in *M. xanthus*. Proc Natl Acad Sci USA 117:23859–23868.

8. Jelsbak L, Kaiser D. 2005. Regulating pilin expression reveals a threshold for S motility in *Myxococcus xanthus*. J Bacteriol 187:2105–2112.

9. Gold VAM, Salzer R, Averhoff B, Kühlbrandt W. 2015. Structure of a type IV pilus machinery in the open and closed state. eLife 4:e07380.

10. Chang Y-W, Rettberg LA, Treuner-Lange A, Iwasa J, Søgaard-Andersen L, Jensen GJ. 2016. Architecture of the type IVa pilus machine. Science 351:aad2001.

11. Treuner-Lange A, Chang Y-W, Glatter T, Herfurth M, Lindow S, Chreifi G, Jensen GJ, Søgaard-Andersen L. 2020. PilY1 and minor pilins form a complex priming the type IVa pilus in *Myxococcus xanthus*. Nat Commun 11:5054.

12. Mancl JM, Black WP, Robinson H, Yang Z, Schubot FD. 2016. Crystal structure of a type IV pilus assembly ATPase: Insights into the molecular mechanism of PilB from *Thermus thermophilus*. Structure 24:1886–1897.

13. Satyshur KA, Worzalla GA, Meyer LS, Heiniger EK, Aukema KG, Misic AM, Forest KT. 2007. Crystal structures of the pilus retraction motor PilT suggest large domain movements and subunit cooperation drive motility. Structure 15:363–376.

14. Misic AM, Satyshur KA, Forest KT. 2010. *P. aeruginosa* PilT structures with and without nucleotide reveal a dynamic type IV pilus retraction motor. J Mol Biol 400:1011–1021.

15. McCallum M, Tammam S, Khan A, Burrows LL, Howell PL. 2017. The molecular mechanism of the type IVa pilus motors. Nat Commun 8:15091.

16. Rudel T, Scheuerpflug I, Meyer TF. 1995. *Neisseria* PilC protein identified as type-4 pilus tip-located adhesin. Nature 373:357–359.

17. Xue S, Mercier R, Guiseppi A, Kosta A, De Cegli R, Gagnot S, Mignot T, Mauriello EMF. 2022. The differential expression of PilY1 proteins by the HsfBA phosphorelay allows twitching motility in the absence of exopolysaccharides. PLOS Genetics 18:e1010188.

18. Herfurth M, Treuner-Lange A, Glatter T, Wittmaack N, Hoiczyk E, Pierik AJ, Søgaard-Andersen L. 2022. A noncanonical cytochrome *c* stimulates calcium binding by PilY1 for type IVa pili formation. Proc Natl Acad Sci U S A 119:e2115061119.

19. Finlay BB, Pasloske BL, Paranchych W. 1986. Expression of the *Pseudomonas aeruginosa* PAK pilin gene in *Escherichia coli*. J Bacteriol 165:625–630.

20. Dupuy B, Taha MK, Pugsley AP, Marchal C. 1991. *Neisseria gonorrhoeae* prepilin export studied in *Escherichia coli*. J Bacteriol 173:7589–7598.

21. Morand PC, Bille E, Morelle S, Eugène E, Beretti JL, Wolfgang M, Meyer TF, Koomey M, Nassif X. 2004. Type IV pilus retraction in pathogenic *Neisseria* is regulated by the PilC proteins. EMBO J 23:2009–2017.

22. Alm R, Bodero A, Free P, Mattick J. 1996. Identification of a novel gene, *pilZ*, essential for type 4 fimbrial biogenesis in Pseudomonas aeruginosa. J Bacteriol 178:46–53.

23. Guzzo CR, Salinas RK, Andrade MO, Farah CS. 2009. PilZ protein structure and interactions with PilB and the FimX EAL domain: Implications for control of type IV pilus biogenesis. J Mol Biol 393: 848–866.

24. Huang B, Whitchurch CB, Mattick JS. 2003. FimX, a multidomain protein connecting environmental signals to twitching motility in *Pseudomonas aeruginosa*. J Bacteriol 185:7068–7076.

25. Jain R, Sliusarenko O, Kazmierczak BI. 2017. Interaction of the cyclic-di-GMP binding protein FimX and the Type 4 pilus assembly ATPase promotes pilus assembly. PLOS Pathogens 13:e1006594.

26. Kazmierczak BI, Lebron MB, Murray TS. 2006. Analysis of FimX, a phosphodiesterase that governs twitching motility in *Pseudomonas aeruginosa*. Mol MIcrobiol 60:1026–1043.

27. Laventie B-J, Sangermani M, Estermann F, Manfredi P, Planes R, Hug I, Jaeger T, Meunier E, Broz P, Jenal U. 2019. A surface-induced asymmetric program promotes tissue colonization by *Pseudomonas aeruginosa*. Cell Host Microbe 25:140–152.e6.

28. Milner DS, Till R, Cadby I, Lovering AL, Basford SM, Saxon EB, Liddell S, Williams LE, Sockett RE. 2014. Ras GTPase-like protein MglA, a controller of bacterial social-motility in Myxobacteria, has evolved to control bacterial predation by *Bdellovibrio*. PLOS Genetics 10:e1004253.

29. Salzer R, Joos F, Averhoff B. 2015. Different effects of MglA and MglB on pilus-mediated functions and natural competence in *Thermus thermophilus*. Extremophiles 19:261–267.

30. Zhang Y, Ducret A, Shaevitz J, Mignot T. 2012. From individual cell motility to collective behaviors: insights from a prokaryote, *Myxococcus xanthus*. FEMS Microbiol Rev 36:149–164.

31. Schumacher D, Søgaard-Andersen L. 2017. Regulation of cell polarity in motility and cell division in *Myxococcus xanthus*. Annu Rev Microbiol 71:61–78.

32. Konovalova A, Petters T, Søgaard-Andersen L. 2010. Extracellular biology of *Myxococcus xanthus*. FEMS Microbiol Rev 34:89–106.

33. Nudleman E, Wall D, Kaiser D. 2006. Polar assembly of the type IV pilus secretin in *Myxococcus xanthus*. Mol Microbiol 60:16–29.

34. Bulyha I, Schmidt C, Lenz P, Jakovljevic V, Höne A, Maier B, Hoppert M, Søgaard-Andersen L. 2009. Regulation of the type IV pili molecular machine by dynamic localization of two motor proteins. Mol Microbiol 74:691–706.

35. Friedrich C, Bulyha I, Søgaard-Andersen L. 2014. Outside-in assembly pathway of the type IV pili system in *Myxococcus xanthus*. J Bacteriol 196:378–390.

36. Siewering K, Jain S, Friedrich C, Webber-Birungi MT, Semchonok DA, Binzen I, Wagner A, Huntley S, Kahnt J, Klingl A, Boekema EJ, Søgaard-Andersen L, van der Does C. 2014. Peptidoglycan-binding protein TsaP functions in surface assembly of type IV pili. Proc Natl Acad Sci USA 111:E953–961.

37. Herfurth M, Pérez-Burgos M, Søgaard-Andersen L. 2023. The mechanism for polar localization of the type IVa pilus machine in *Myxococcus xanthus*. mBio 14:e01593–23.

38. Kaiser D. 1979. Social gliding is correlated with the presence of pili in *Myxococcus xanthus*. Proc Natl Acad Sci USA 76:5952–5956.

39. Sun H, Zusman DR, Shi W. 2000. Type IV pilus of *Myxococcus xanthus* is a motility apparatus controlled by the *frz* chemosensory system. Curr Biol 10:1143–1146.

40. Blackhart BD, Zusman DR. 1985. “Frizzy” genes of *Myxococcus xanthus* are involved in control of frequency of reversal of gliding motility. Proc Natl Acad Sci USA 82:8771–8774.

41. Carreira LAM, Szadkowski D, Müller F, Søgaard-Andersen L. 2022. Spatiotemporal regulation of switching front–rear cell polarity. Curr Opin Cell Biol 76:102076.

42. Dinet C, Mignot T. 2023. Unorthodox regulation of the MglA Ras-like GTPase controlling polarity in *Myxococcus xanthus*. FEBS Letters 597:850–864.

43. Carreira LAM, Szadkowski D, Lometto S, Hochberg GKA, Søgaard-Andersen L. 2023. Molecular basis and design principles of switchable front-rear polarity and directional migration in *Myxococcus xanthus*. Nat Commun 14:4056.

44. Leonardy S, Miertzschke M, Bulyha I, Sperling E, Wittinghofer A, Søgaard-Andersen L. 2010. Regulation of dynamic polarity switching in bacteria by a Ras-like G-protein and its cognate GAP. EMBO J 29:2276–2289.

45. Zhang Y, Franco M, Ducret A, Mignot T. 2010. A bacterial Ras-like small GTP-binding protein and its cognate GAP establish a dynamic spatial polarity axis to control directed motility. PLOS Biol 8:e1000430.

46. Mercier R, Bautista S, Delannoy M, Gibert M, Guiseppi A, Herrou J, Mauriello EMF, Mignot T. 2020. The polar Ras-like GTPase MglA activates type IV pilus via SgmX to enable twitching motility in *Myxococcus xanthus*. Proc Natl Acad Sci USA 117:28366–28373.

47. Hodgkin J, Kaiser D. 1979. Genetics of gliding motility in *Myxococcus xanthus* (Myxobacterales): Two gene systems control movement. Mol Gen Genet 171:177–191.

48. Hartzell P, Kaiser D. 1991. Function of MglA, a 22-kilodalton protein essential for gliding in *Myxococcus xanthus*. J Bacteriol 173:7615–24.

49. Szadkowski D, Harms A, Carreira LAM, Wigbers M, Potapova A, Wuichet K, Keilberg D, Gerland U, Søgaard-Andersen L. 2019. Spatial control of the GTPase MglA by localized RomR/RomX GEF and MglB GAP activities enables *Myxococcus xanthus* motility. Nat Microbiol 4:1344–1355.

50. Szadkowski D, Carreira LAM, Søgaard-Andersen L. 2022. A bipartite, low-affinity roadblock domain-containing GAP complex regulates bacterial front-rear polarity. PLOS Genetics 18:e1010384.

51. Carreira LAM, Tostevin F, Gerland U, Søgaard-Andersen L. 2020. Protein-protein interaction network controlling establishment and maintenance of switchable cell polarity. PLOS Genet 16:e1008877.

52. Leonardy S, Freymark G, Hebener S, Ellehauge E, Søgaard-Andersen L. 2007. Coupling of protein localization and cell movements by a dynamically localized response regulator in *Myxococcus xanthus*. EMBO J 26:4433–4444.

53. Zhang Y, Guzzo M, Ducret A, Li Y-Z, Mignot T. 2012. A dynamic response regulator protein modulates G-protein–dependent polarity in the bacterium *Myxococcus xanthus*. PLOS Genet 8:e1002872.

54. Keilberg D, Wuichet K, Drescher F, Søgaard-Andersen L. 2012. A response regulator interfaces between the Frz chemosensory system and the MglA/MglB GTPase/GAP module to regulate polarity in *Myxococcus xanthus*. PLOS Genet 8:e1002951.

55. Mauriello EMF, Mouhamar F, Nan B, Ducret A, Dai D, Zusman DR, Mignot T. 2010. Bacterial motility complexes require the actin-like protein, MreB and the Ras homologue, MglA. EMBO J 29:315–326.

56. Fraser JS, Merlie Jr JP, Echols N, Weisfield SR, Mignot T, Wemmer DE, Zusman DR, Alber T. 2007. An atypical receiver domain controls the dynamic polar localization of the *Myxococcus xanthus* social motility protein FrzS. Mol Microbiol 65:319–332.

57. Ward MJ, Lew H, Zusman DR. 2000. Social motility in *Myxococcus xanthus* requires FrzS, a protein with an extensive coiled-coil domain. Mol Microbiol 37:1357–1371.

58. Bautista S, Schmidt V, Guiseppi A, Mauriello EMF, Attia B, Elantak L, Mignot T, Mercier R. 2023. FrzS acts as a polar beacon to recruit SgmX, a central activator of type IV pili during *Myxococcus xanthus* motility. EMBO J 42:e111661.

59. Szklarczyk D, Kirsch R, Koutrouli M, Nastou K, Mehryary F, Hachilif R, Gable AL, Fang T, Doncheva NT, Pyysalo S, Bork P, Jensen LJ, von Mering C. 2022. The STRING database in 2023: protein–protein association networks and functional enrichment analyses for any sequenced genome of interest. Nucleic Acids Res 51:D638–D646.

60. Potapova A. 2019. Regulation of type IV pili formation and function by the small GTPase MglA in *Myxococcus xanthus*. PhD thesis. Philipps-Universität Marburg.

61. Kuzmich S, Blumenkamp P, Meier D, Szadkowski D, Goesmann A, Becker A, Søgaard-Andersen L. 2022. CRP-like transcriptional regulator MrpC curbs c-di-GMP and 3’,3’-cGAMP nucleotide levels during development in *Myxococcus xanthus*. mBio 13:e00044–22.

62. Shi W, Zusman DR. 1993. The two motility systems of *Myxococcus xanthus* show different selective advantages on various surfaces. Proc Natl Acad Sci USA 90:3378–3382.

63. Wu SS, Kaiser D. 1995. Genetic and functional evidence that type IV pili are required for social gliding motility in *Myxococcus xanthus*. Mol Microbiol 18:547–558.

64. Nan B, Bandaria JN, Moghtaderi A, Sun I-H, Yildiz A, Zusman DR. 2013. Flagella stator homologs function as motors for myxobacterial gliding motility by moving in helical trajectories. Proc Natl Acad Sci USA 110:E1508–E1513.

65. Sun M, Wartel M, Cascales E, Shaevitz JW, Mignot T. 2011. Motor-driven intracellular transport powers bacterial gliding motility. Proc Natl Acad Sci USA 108:7559–7564.

66. Jakovljevic V, Leonardy S, Hoppert M, Søgaard-Andersen L. 2008. PilB and PilT are ATPases acting antagonistically in type IV pili function in *Myxococcus xanthus*. J Bacteriol 190:2411–2421.

67. Wu SS, Wu J, Kaiser D. 1997. The *Myxococcus xanthus pilT* locus is required for social gliding motility although pili are still produced. Mol Microbiol 23:109–121.

68. Mignot T, Merlie JP, Zusman DR. 2005. Regulated pole-to-pole oscillations of a bacterial gliding motility protein. Science 310:855–857.

69. Karimova G, Pidoux J, Ullmann A, Ladant D. 1998. A bacterial two-hybrid system based on a reconstituted signal transduction pathway. Proc Natl Acad Sci USA 95:5752–5756.

70. Mignot T, Merlie Jr JP, Zusman DR. 2007. Two localization motifs mediate polar residence of FrzS during cell movement and reversals of *Myxococcus xanthus*. Mol Microbiol 65:363–372.

71. Bischof LF, Friedrich C, Harms A, Søgaard-Andersen L, van der Does C. 2016. The type IV pilus assembly ATPase PilB of *Myxococcus xanthus* interacts with the inner membrane platform protein PilC and the nucleotide-binding protein PilM. J Biol Chem 291:6946–6957.

72. McCallum M, Tammam S, Little DJ, Robinson H, Koo J, Shah M, Calmettes C, Moraes TF, Burrows LL, Howell PL. 2016. PilN binding modulates the structure and binding partners of the *Pseudomonas aeruginosa* type IVa pilus protein PilM. J Biol Chem 291:11003–11015.

73. Jones CJ, Utada A, Davis KR, Thongsomboon W, Zamorano Sanchez D, Banakar V, Cegelski L, Wong GCL, Yildiz FH. 2015. C-di-GMP regulates motile to sessile transition by modulating MshA pili biogenesis and near-surface motility behavior in *Vibrio cholerae*. PLOS Pathogens 11:e1005068.

74. Roelofs KG, Jones CJ, Helman SR, Shang X, Orr MW, Goodson JR, Galperin MY, Yildiz FH, Lee VT. 2015. Systematic identification of cyclic-di-GMP binding proteins in *Vibrio cholerae* reveals a novel class of cyclic-di-GMP-binding ATPases associated with type II secretion systems. PLOS Pathogens 11:e1005232.

75. Hendrick WA, Orr MW, Murray SR, Lee VT, Melville SB. 2017. Cyclic Di-GMP binding by an assembly ATPase (PilB2) and control of type IV pilin polymerization in the Gram-positive pathogen *Clostridium perfringens*. J Bacteriol 199:e00034–17.

76. Guzzo CR, Dunger G, Salinas RK, Farah CS. 2013. Structure of the PilZ–FimXEAL– c-di-GMP complex responsible for the regulation of bacterial type IV pilus biogenesis. J Mol Biol 425:2174–2197.

77. Llontop EE, Cenens W, Favaro DC, Sgro GG, Salinas RK, Guzzo CR, Farah CS. 2021. The PilB-PilZ-FimX regulatory complex of the Type IV pilus from *Xanthomonas citri*. PLOS Pathogens 17:e1009808.

78. Pérez-Burgos M, García-Romero I, Jung J, Schander E, Valvano MA, Søgaard-Andersen L. 2020. Characterization of the exopolysaccharide biosynthesis pathway in *Myxococcus xanthus*. J Bacteriol 202:e00335–20.

79. Schwabe J, Pérez-Burgos M, Herfurth M, Glatter T, Søgaard-Andersen L. 2022. Evidence for a widespread third system for bacterial polysaccharide export across the outer membrane comprising a composite OPX/β-barrel translocon. mBio 13:e02032–22.

80. Hodgkin J, Kaiser D. 1977. Cell-to-cell stimulation of movement in nonmotile mutants of *Myxococcus*. Proc Natl Acad Sci USA 74:2938–2942.

81. Shi X, Wegener-Feldbrügge S, Huntley S, Hamann N, Hedderich R, Søgaard-Andersen L. 2008. Bioinformatics and experimental analysis of proteins of two-component systems in *Myxococcus xanthus*. J Bacteriol 190:613–624.

82. Sambrook J, Russell DW. 2001. Molecular Cloning: A Laboratory Manual, 3rd ed. Cold Spring Harbor Laboratory Press, Cold Spring Harbor, N.Y.

83. Schneider CA, Rasband WS, Eliceiri KW. 2012. NIH Image to ImageJ: 25 years of image analysis. Nat Methods 9:671–675.

84. Schindelin J, Arganda-Carreras I, Frise E, Kaynig V, Longair M, Pietzsch T, Preibisch S, Rueden C, Saalfeld S, Schmid B, Tinevez JY, White DJ, Hartenstein V, Eliceiri K, Tomancak P, Cardona A. 2012. Fiji: an open-source platform for biological-image analysis. Nat Methods 9:676–82.

85. Paintdakhi A, Parry B, Campos M, Irnov I, Elf J, Surovtsev I, Jacobs-Wagner C. 2016. Oufti: an integrated software package for high-accuracy, high-throughput quantitative microscopy analysis. Mol Microbiol 99:767–777.

86. Madeira F, Pearce M, Tivey ARN, Basutkar P, Lee J, Edbali O, Madhusoodanan N, Kolesnikov A, Lopez R. 2022. Search and sequence analysis tools services from EMBL-EBI in 2022. Nucl Acids Res 50:W276–W279.

87. Waterhouse AM, Procter JB, Martin DMA, Clamp M, Barton GJ. 2009. Jalview Version 2—a multiple sequence alignment editor and analysis workbench. Bioinformatics 25:1189–1191.

88. Blum M, Chang H-Y, Chuguransky S, Grego T, Kandasaamy S, Mitchell A, Nuka G, Paysan-Lafosse T, Qureshi M, Raj S, Richardson L, Salazar GA, Williams L, Bork P, Bridge A, Gough J, Haft DH, Letunic I, Marchler-Bauer A, Mi H, Natale DA, Necci M, Orengo CA, Pandurangan AP, Rivoire C, Sigrist CJA, Sillitoe I, Thanki N, Thomas PD, Tosatto SCE, Wu CH, Bateman A, Finn RD. 2020. The InterPro protein families and domains database: 20 years on. Nucl Acids Res 49:D344–D354.

89. Aoki KF, Kanehisa M. 2005. Using the KEGG database resource. Curr Protoc Bioinformatics 11:1.12.1–1.12.54.

90. Li W, Cowley A, Uludag M, Gur T, McWilliam H, Squizzato S, Park YM, Buso N, Lopez R. 2015. The EMBL-EBI bioinformatics web and programmatic tools framework. Nucl Acids Res 43:W580–W584.

91. Kumar S, Stecher G, Tamura K. 2016. MEGA7: Molecular evolutionary genetics analysis Version 7.0 for bigger datasets. Mol Biol Evol 33:1870–1874.

92. Wu SS, Kaiser D. 1996. Markerless deletions of *pil* genes in *Myxococcus xanthus* generated by counterselection with the *Bacillus subtilis sacB* gene. J Bacteriol 178:5817–5821.

93. Jakobczak B, Keilberg D, Wuichet K, Søgaard-Andersen L. 2015. Contact- and protein transfer-dependent stimulation of assembly of the gliding motility machinery in *Myxococcus xanthus*. PLOS Genet 11:e1005341.

94. Julien B, Kaiser AD, Garza A. 2000. Spatial control of cell differentiation in *Myxococcus xanthus*. Proc Natl Acad Sci U S A 97:9098–9103.

95. Miertzschke M, Körner C, Vetter IR, Keilberg D, Hot E, Leonardy S, Søgaard-Andersen L, Wittinghofer A. 2011. Structural analysis of the Ras-like G protein MglA and its cognate GAP MglB and implications for bacterial polarity. EMBO J 30:4185–4197.

