## Supplementary Information for "Combinatorial control of type IVa pili formation by the four polarized regulators MglA, SgmX, FrzS and SopA"

**This file contains:**

- Supplementary Figures 1-7
- Supplementary Table 1-2

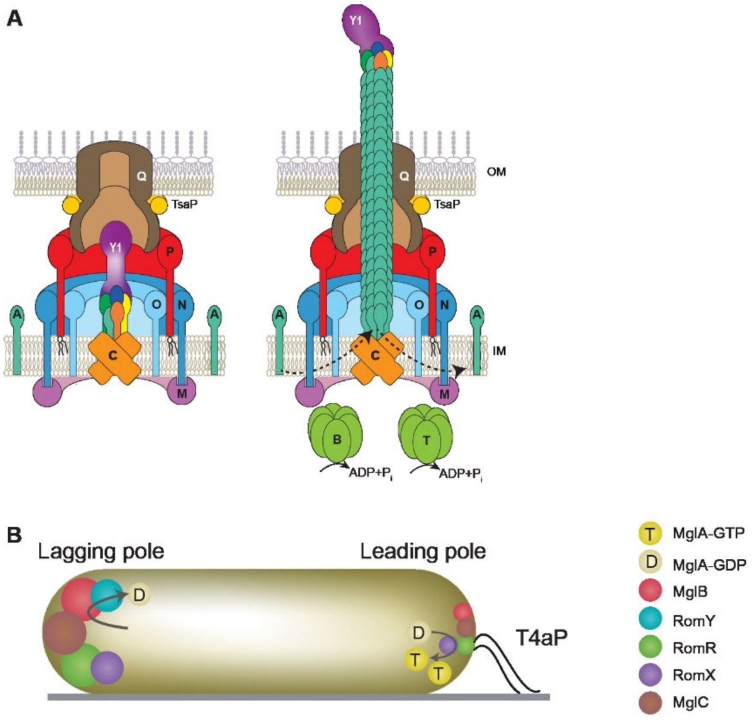


**Supplementary Figure 1.** Schematics of the T4aPM and the proteins of the polarity module. **A.** Architectural model of T4aPM in two states with the 15 conserved proteins. Left, unpiliated T4aPM. The core T4aPM consists of four functional elements, i.e. an OM pore (PilQ and TsaP), an alignment complex (PilP, PilN and PilO) that connects the OM pore to the IM platform complex (PilC and PilM), and a priming complex (four minor pilins (orange, yellow, blue and green) and PilY1). Right, piliated T4aPM. As in the left panel except that the T4aPM contains an extended T4aP capped by the tip complex. PilB and PilT associate with PilM and PilC in a mutually exclusive manner for extension and retraction, respectively. Bent arrows indicate incorporation at and removal from the pilus base of PilA during extension and retraction, respectively. Proteins labeled with single letters have the Pil prefix. **B.** Localization of proteins of the polarity module. T4aP are shown at the leading pole. Bent arrows at the leading and lagging poles indicate that the RomR/RomR GEF stimulates the exchange of GDP for GTP in MglA at the leading pole and the MglB/RomY complex stimulates the low intrinsic GTPase activity of MglA at the lagging pole. The size of circles indicates the relative amount of a protein at a pole.

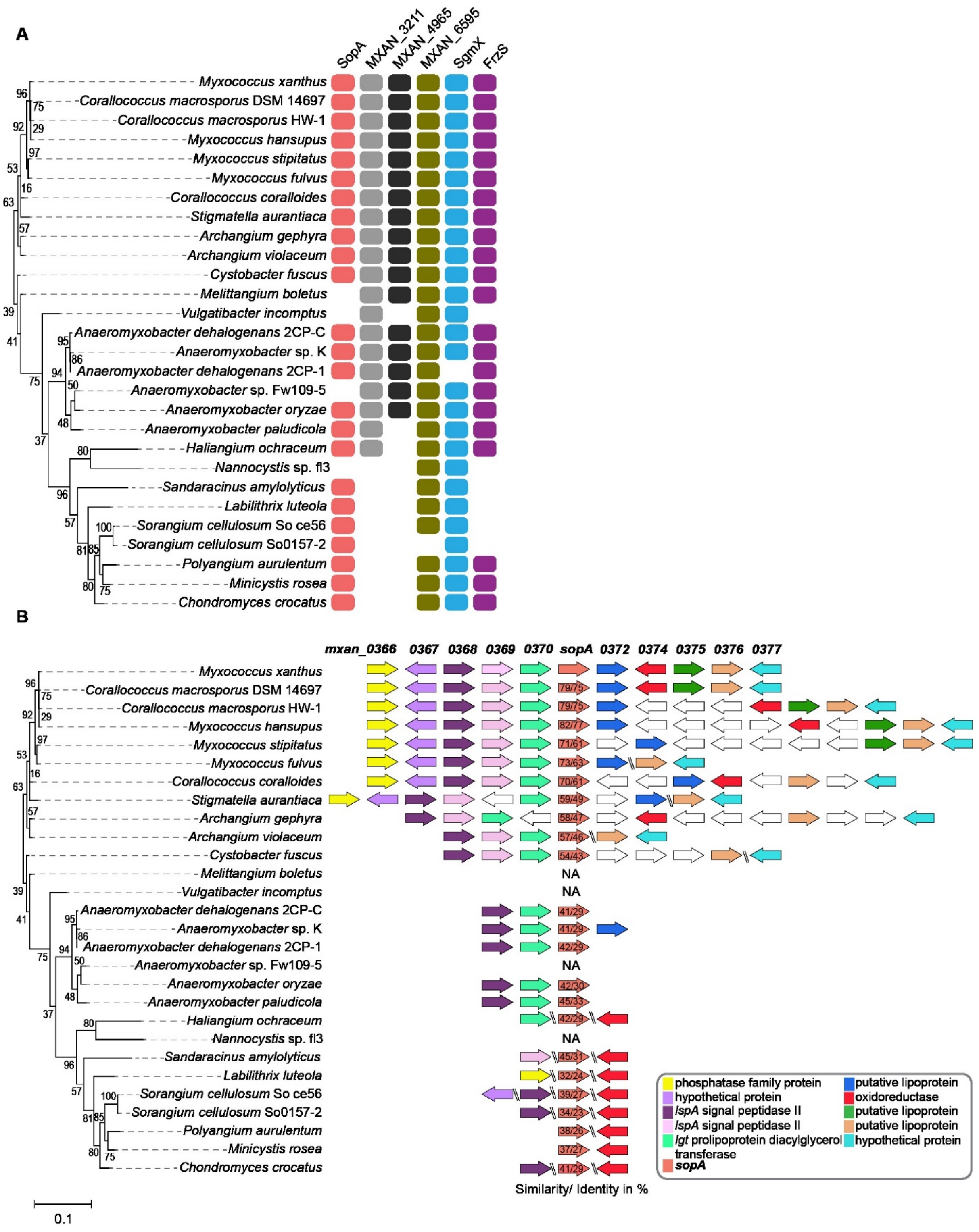


**Supplementary Figure 2**. SopA co-occurs with SgmX in Myxococcales and the *sopA* locus is conserved in related Myxococcales.

**A.** Occurrence of SopA, SgmX, FrzS, MXAN_3211, MXAN_4965 and MXAN_6595 in Myxococcales with fully sequenced genomes. Orthologs were identified using the KEGG SSDB database. **B.** The *sopA* locus is conserved in related Myxoccocales. Transcription direction is indicated by the orientation of arrows with MXAN numbers indicated for the *sopA* locus in *M. xanthus*. % similarity/identity between SopA homologs from *M. xanthus* and other species is indicated by numbers in the arrows. For the proteins encoded by genes flanking *sopA* in *M. xanthus*, domains were identified using Interpro. % similarity/identity between protein homologs were calculated using EMBOSS Needle software (pairwise sequence alignment). In A and B, phylogenetic trees were prepared in MEGA7 using the Neighbor-Joining method. Bootstrap values (500 replicates) are shown next to the branches.

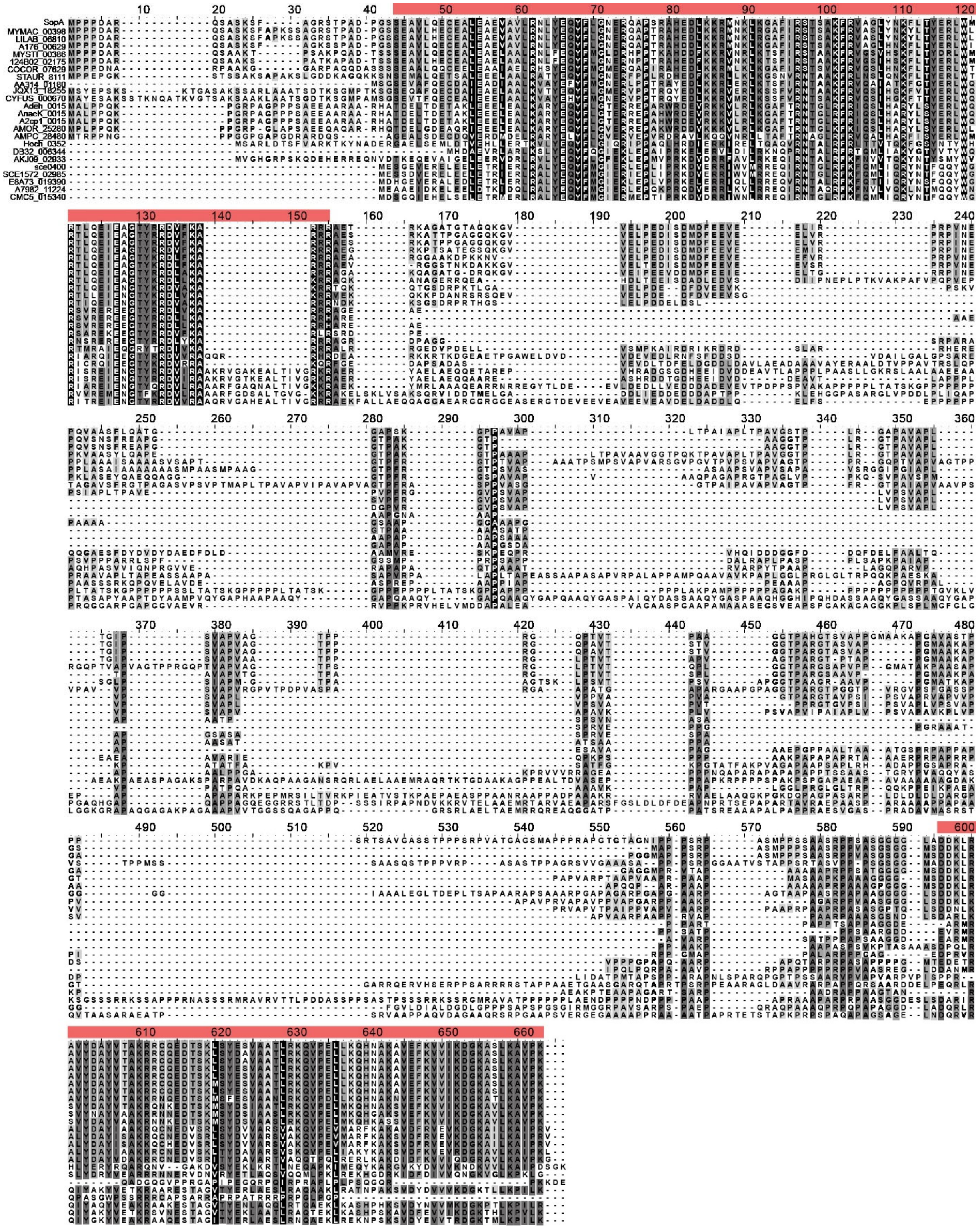


**Supplementary Figure 3.** Sequence alignment of SopA homologs. Light red bars indicate the conserved N-terminal and C-terminal regions.

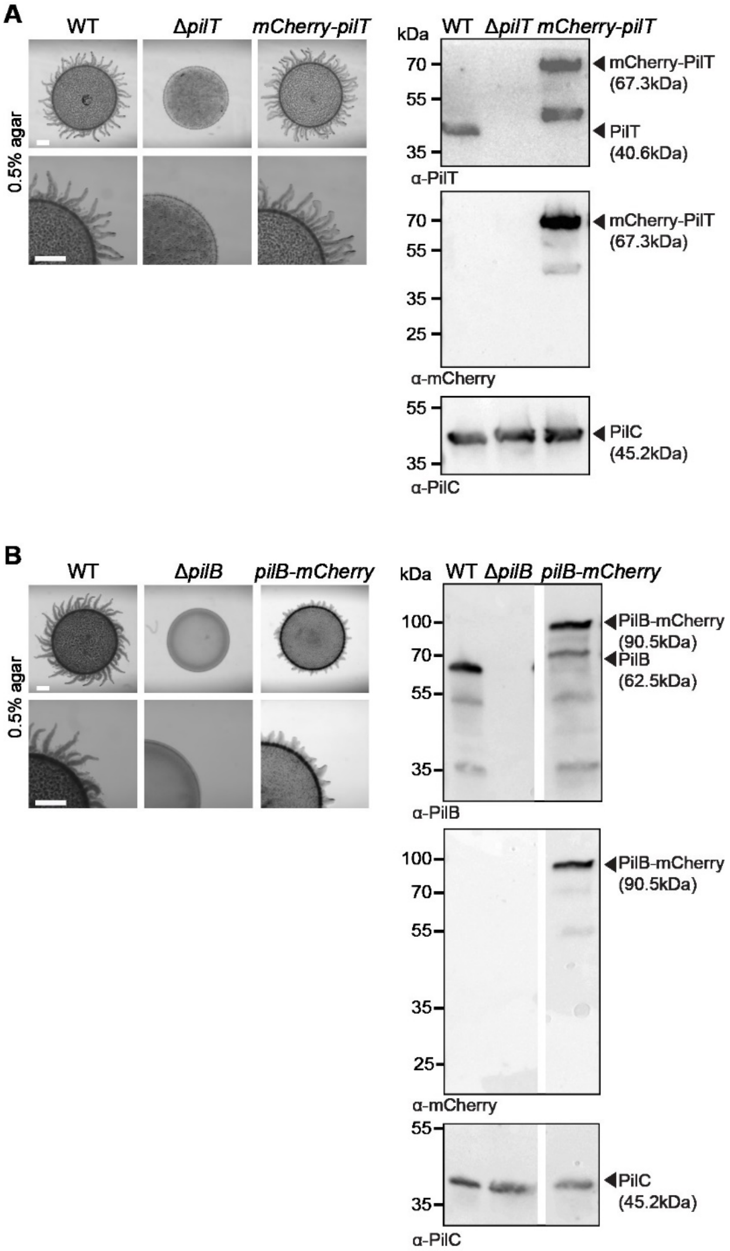


**Supplementary Figure 4.** Analysis of strains synthesizing mCherry-PilT or PilB-mCherry.
**A, B.** The mCherry-PilT is fully active and the PilB-mCherry fusion partially active. Left panels, cells were incubated on 0.5% agar supplemented with 0.5% CTT to score T4aP-dependent. Images were recorded at 24 h. Scale bars, 1mm. Both fusion proteins were synthesized from their native locus. Right panels, accumulation of mCherry-PilT and PilB-mCherry assessed by immunoblotting. Protein from total cell extracts of $10^8$ cells was separated by SDS-PAGE and probed with α-PilT/PilB antibodies (top), then after stripping with α-mCherry antibodies (middle), and then after stripping with α-PilC antibodies as a loading control (bottom).

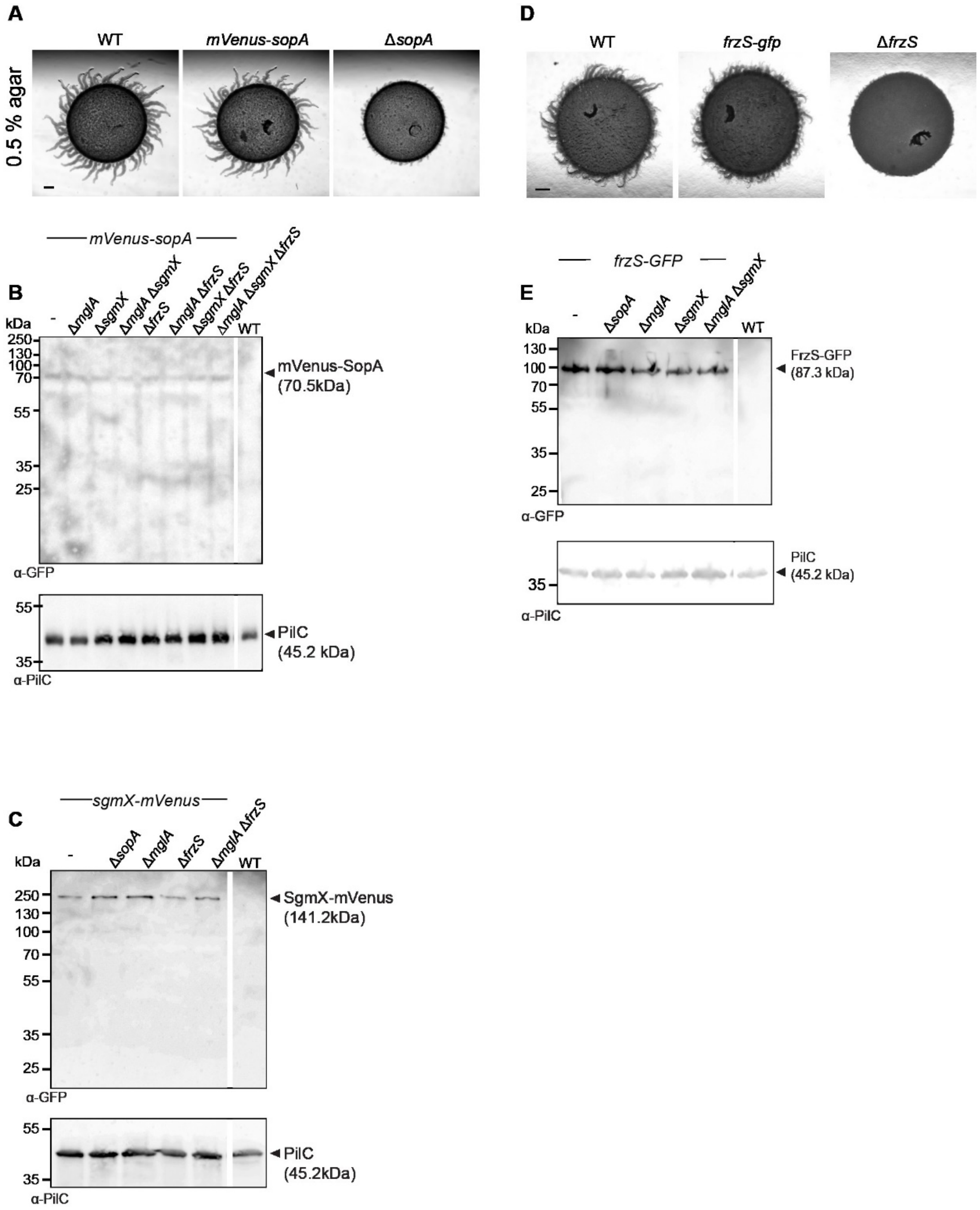


**Supplementary Figure 5.** Analysis of the accumulation of mVenus-SopA, SgmX-mVenus and FrzS-GFP.

**A, D.** The mVenus-SopA (A) and FrzS-GFP (D) fusion proteins are fully active. Cells were incubated on 0.5% agar supplemented with 0.5% CTT to score T4aP-dependent and imaged at 24 h. Scale bars, 1 mm. Both proteins were synthesized from their native locus. **B, C and E**. Immunoblots to assess the accumulation of the indicated fusion proteins. Protein from total cell extracts of $10^8$ cells was separated by SDS-PAGE and probed with the indicated antibodies (top), and then after stripping with α-PilC antibodies as a loading control (bottom). Gaps indicate lanes removed for presentation purposes.

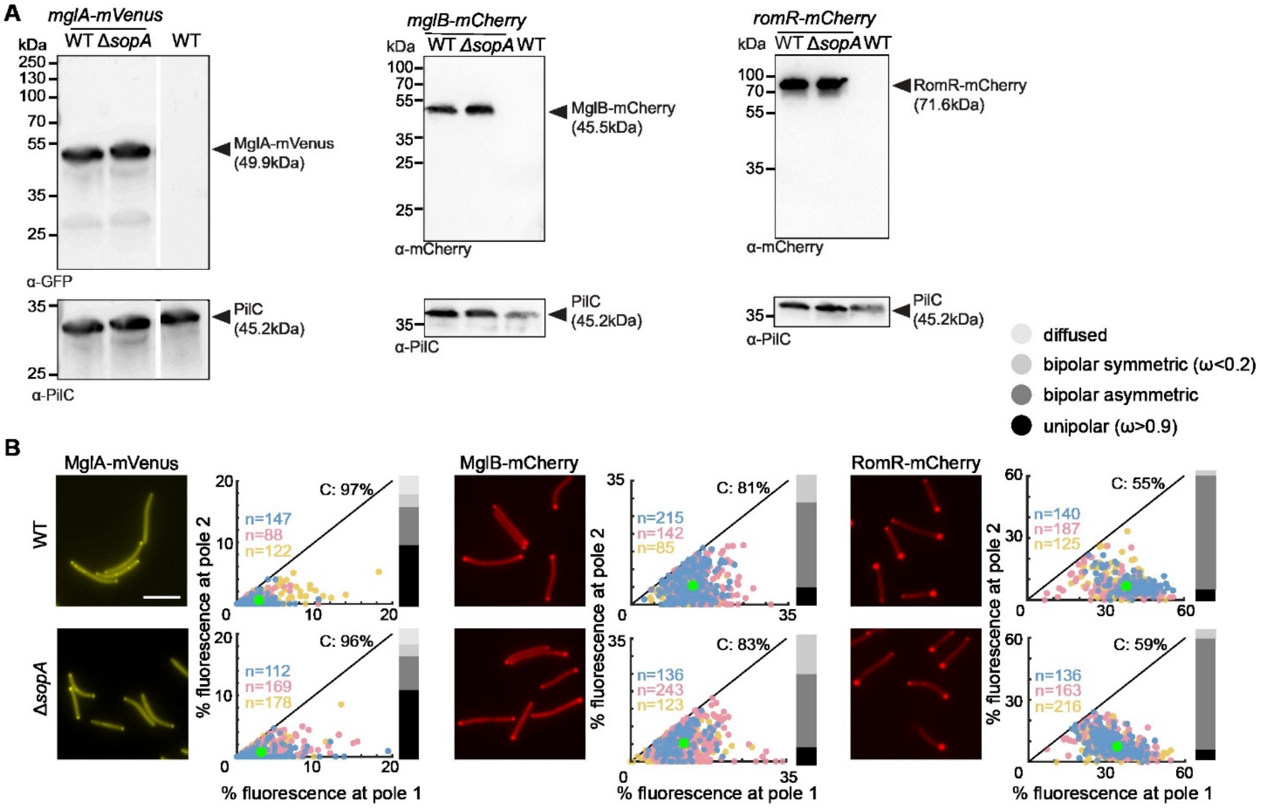


**Figure 6.** SopA is neither important for polar MglA-mVenus, MglB-mCherry and RomR-mCherry accumulation nor polar localization.

**A.** Immunoblots to assess the accumulation of the indicated fusion proteins. Protein from total cell extracts of $10^8$ cells was separated by SDS-PAGE and probed with the indicated antibodies (top), and then after stripping with α-PilC antibodies as a loading control (bottom). Gap indicates lanes removed for presentation purposes. **B.** Experiments were done, presented and analyzed as in Fig. 2D.

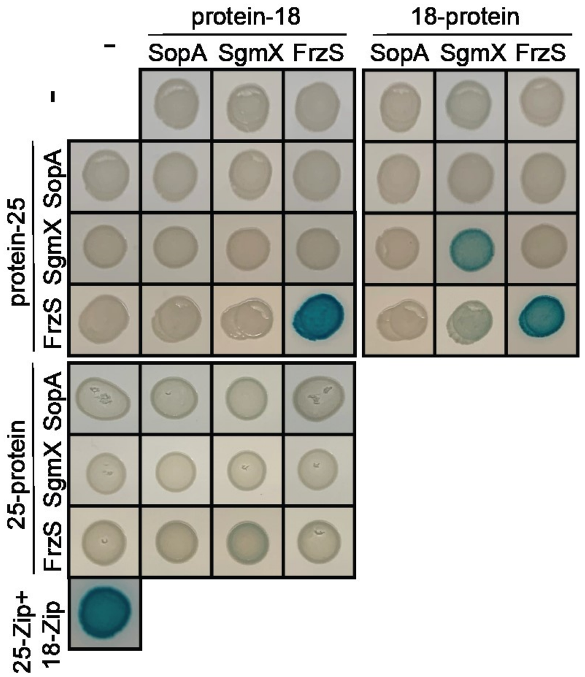


**Supplementary Figure 7.** BACTH assay for SgmX, FrzS and SopA interactions.
Full-length SgmX, FrzS and SopA were fused to the N-terminus or C-terminus of T25 and T18.
Lower left corner, T25-Zip + T18-Zip positive control.

**Table S1**. Proteins co-occurring with SgmX based on STRING search

| **Locus tag (old)** | **Locus tag (new)** | **Description** |
|---|---|---|
| MXAN_0371 (SopA) | MXAN_RS01825 | Hypothetical protein |
| MXAN_0869 | MXAN_RS04165 | Triacylglycerol lipase |
| MXAN_1397 | MXAN_RS06775 | Phycobilisome lyase |
| MXAN_2696 | MXAN_RS13060 | Desaturase |
| MXAN_3211 | MXAN_RS15550 | PATAN domain protein |
| MXAN_4965 | MXAN_RS24110 | PATAN domain protein |
| MXAN_6595 | MXAN_RS24110 | TPR domain protein |
| MXAN_5763 | MXAN_RS27935 | Omp85 domain protein |
| MXAN_5764 | MXAN_RS27940 | TamB-like protein |
| MXAN_5765 | MXAN_RS27945 | SecA-like protein |

**Table S2.** Primers used in this work

| Primer name | Sequence | |
|---|---|---|
| *mxan0371_A* | ATCGAAGCTTGTGGAAGCTACGGGTGAAAT | Primers for generation of *sopA* in-frame deletion |
| *mxan0371_B* | GAGCGATGCGGACTGTCGGGCGTCGGG | |
| *mxan0371_C* | CGACAGTCCGCATCGCTCAAGGCCGTG | |
| *mxan0371_D* | ATCGGAATTCCATCATCGCGATCATGTGTG | |
| *mxan0371_E* | ACACGTCCAGCCTGGCGTATT | Primers for checking *sopA* in-frame deletion, and *mVenus-sopA* fusion |
| *mxan0371_F* | CCGCGCCGCTTGAGCTG | |
| *mxan0371_G* | GAAATCGAAGCGGGCACCTATCGCCGG | |
| *mxan0371_H* | CGGTGACGTACGCGTCGTAGACGGCAC | |
| 0371 nat. promotor AHind3 | GCGCAAGCTTGTCGTGGAAGCTACGGGT | Primers for generation of the complementation plasmid |
| Mxan_0371 B EcoRI (Van) | GCGCGAATTCCTACTTCGGCACGGCCTT | |
| mVenus-0371 fus. A Hind3 | GCGCAAGCTTGTGTTCTCCCTGGGCGGA | Primers for generation of *mVenus-sopA* fusion |
| mVenus amp. forward | GGGACACTCTGAGGAGTCATGCTGAGCAAG | |
| mVenus-0371 fusion B | GCCCTTGCTCACCATCACTCCTCAGAGTGT | |
| mVenus amp. rev + linker 1 | GGATCCTCCTCCTCCGGAGCCGCCGCCGCCCTTG TACAGCTCGTCCAT | |
| mVenus-0371 C1 | GGCGGCGGCGGCTCCGGAGGAGGAGGATCCATG CCGCCCCCCGACGCC | |
| mVenus-0371 D EcoRI 2 | GCGCGAATTCGGTGCCTTCGCCGCCATGCC | |
| FrzS_E new | ACGAGTGGACCTCGAAACCCACC | Primers for checking *frzS* in-frame deletion, and *frzS-gfp* |
| FrzS_F | CACGTTCGACCCGGACGCGAA | |
| FrzS_G | GGGGCTTCACGGTCGACG | |
| FrzS_H | TTCTTGTTGGCCGCGTCGC | |
| PilB E2 | CAGGCAAGGTGCTCCAGCCG | Primers for checking *pilB* in-frame deletion, and *pilB-mCherry* fusion |
| PilB 1 F | GCGTCGCGTAGCAGATGTG | |
| PilB 1 G | GCTTCGACGCGCAGCCGCTG | |
| PilB 1 H | GGCCACGCGGCCGCGGTAGC | |
| pilT-E | CTCCGCCAGGACCCGGACATC | Primers for checking *pilT* in-frame deletion, and *mCherry-pilT* fusion |
| pilT-F | CGAAGACGGGCGTCACCTTC | |
| 5787-G pilT | CTTGAAGACGGCGCCGCTGA | |
| 5787-H pilT | CGCGCTGATTCACGAGGCAG | |
| B2H Mxan_0371 fw XbaI new | GCGCTCTAGATATGCCGCCCCCGACGCC | Primer for *sopA* BACTH constructs |
| B2H Mxan_371 revBamHInew | GCGCGGATCCACCTTCGGCACGGCCTTGAG | |
| FrzS B2H rev EcoRI new | GCGCGAATTCGTGGCCGCGGCTTCGCTGGC | Primer for *frzS* BACTH constructs |
| FrzS B2H fwd XbaI new | GCGCTCTAGATATGTCGAAGAAAATCCTG | |
